# BC-Design: A Biochemistry-Aware Framework for Inverse Protein Design

**DOI:** 10.1101/2024.10.28.620755

**Authors:** Xiangru Tang, Xinwu Ye, Fang Wu, Yimeng Liu, Anna Su, Antonia Panescu, Guanlue Li, Daniel Shao, Dong Xu, Mark Gerstein

## Abstract

Inverse protein design (IPD) extends the classical problem of inverse protein folding (IPF) by not only recovering a sequence compatible with a given backbone geometry but also generating a range of new sequences that satisfy the physicochemical constraints required for a stable fold with a specified (often designed) function. Recent progress has incorporated biochemical properties as discrete per-residue features, but such localized signals cannot represent the continuous hydrophobic and electrostatic environments that span protein surfaces and interior volumes. Here we introduce BC-Design, a biochemistry-aware inverse design framework that integrates geometric structure with smoothly varying hydrophobicity and charge properties. The later are represented independent of residue coordinates, on compact point clouds sampled throughout protein surfaces and interiors. This formulation provides a natural spatial description of physicochemical environments and enriches structural cues without specifying amino-acid identities. On the CATH 4.2 benchmark, BC-Design achieves 90% sequence recovery and generalizes robustly across protein lengths, contact-order regimes, and major fold classes. Masking experiments further demonstrate an interpretable fidelity–diversity control: full biochemical context yields high-fidelity reconstructions, whereas withholding information enables exploratory variant generation. Beyond diagnostic benchmarks, BC-Design demonstrably improves functional design outcomes across diverse settings. It increases enzyme–substrate affinity, enhances peptide–receptor design accuracy, and achieves state-of-the-art recovery and structural fidelity in antibody loop (CDRH3) modeling. In these applications, the model not only reconstructs native-like sequences but also proposes plausible functional variants that preserve catalytic or binding geometry—such as generating CDRH3 loops that maintain antigen-contact configurations while offering new sequence solutions. These results show that integrating continuous physicochemical properties with structural geometry enables practical, function-oriented protein design, going beyond backbone-conditioned recovery.

## 1 Introduction

Inverse protein design (IPD) seeks to determine amino acid sequences that are compatible with a target three-dimensional structure and its underlying physicochemical constraints (see Fig. 1). Recent advances in inverse protein folding (IPF) have shown that backbone geometry provides a strong structural scaffold on which models can aim to infer native-like sequences. Building on this foundation, IPD moves beyond sequence recovery: rather than merely reproducing the wild-type sequence, the goal is to design new sequences that not only maintain the intended fold but also satisfy functional or application-specific constraints, such as improving binding, altering activity, or modulating interaction specificity. Given this expanded scope, IPD has become a central problem in computational biology, underpinning a wide spectrum of modern protein engineering efforts [1–7]. By learning the biophysical determinants of sequence– structure compatibility, IPD enables systematic exploration of fold-accessible sequence space and supports the design of enzymes, binders, and other functional proteins with tailored activities [8–12]. Because many residues tolerate extensive mutation without altering the global fold, the fundamental challenge of IPD lies in modeling the geometric and energetic constraints that define viable protein structures, rather than recovering specific native sequences.

**Fig. 1:**
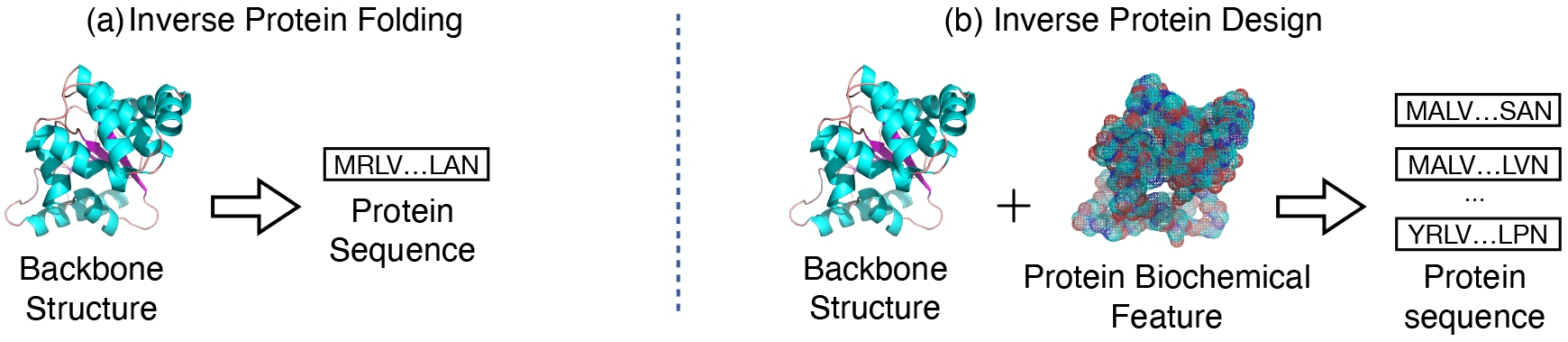
Comparison between traditional inverse protein folding and inverse protein design. (a) Conventional inverse protein folding models take only the backbone structure as input and predict a single amino-acid sequence, treating the protein purely as a geometric object and aiming to recover the native sequence as accurately as possible. (b) Inverse protein design aims to identify structure-compatible sequences that meet specific design objectives, including generating multiple viable sequence solutions or optimizing sequences for improved binding or activity. BC-Design augments backbone geometry with continuous biochemical properties—hydrophobicity and charge—represented over compact point clouds sampled from both protein surfaces and interior regions. This biochemistry-aware formulation enables the model to learn how structural environments and physicochemical gradients jointly shape sequence preferences, supporting both exploratory sequence diversification and targeted function-oriented design. Although biochemical properties enhance learning and yield the best performance, the model remains compatible with backbone-only inference.

A wide range of IPF architectures has been developed [2, 4, 13–22], spanning MLPs, CNNs, graph networks, transformers, and models augmented with residue- or atom-level biochemical annotations. Although these approaches have substantially advanced backbone-conditioned sequence recovery, they treat biochemical information as discrete, index-based labels and therefore cannot capture the smooth energetic properties that govern protein folding. Recent functional IPD frameworks have taken an important step forward by incorporating surface geometry and local chemistry as conditioning signals. SurfPro conditions sequence generation on molecular surfaces with biochemical attributes; DS-ProGen [23] jointly encodes backbone structure and surface chemical geometry within a next-amino-acid prediction framework; and surface-driven generative models such as SurfFlow [24] and PepBridge [25] use flow matching or diffusion bridges to enforce receptor–ligand surface complementarity. SurfDesign [26] further operationalizes surface-aware modeling through equivariant message passing on molecular surfaces combined with PLM fine-tuning, yielding strong gains across CATH, TS50/500, and PDB benchmarks. Existing methods typically encode biochemical information either as residue-centered properties or through point clouds concentrated on the molecular surface (see Appendix 5). While these representations capture local or interfacial chemistry and work well for binder and peptide design, they do not reflect the broader physicochemical environment, including both exterior surfaces and interior cavities, that governs folding stability and global energetics.

Building on these observations, we introduce BC-Design, a biochemistry-aware inverse design framework that models biochemical context across the entire protein rather than only at exposed surfaces or local contact regions. BC-Design distributes hydrophobicity and charge as smoothly varying spatial signals on a compact point-cloud representation that spans both surface and interior regions. These two physicochemical properties capture dominant contributors to protein energetics while remaining sufficiently coarse to avoid revealing residue identities. Ablation analyses further show that enlarging the feature set offers minimal additional benefit and can reduce robustness when biochemical information is absent, indicating that hydrophobicity and charge constitute an efficient and principled biochemical basis for inverse design.

Because biochemical properties may not always be available in practical design settings, BC-Design is constructed to operate under both full-information and backbone-only conditions. Our goal is therefore not to remove biochemical information at inference, but to leverage it during training while retaining the flexibility to function when such properties are absent. While backbone-only inference is fully supported, the model attains its best performance when biochemical context is provided.

Beyond this flexibility, BC-Design provides a practical mechanism for tuning the balance between conservative and exploratory sequence design. During training, stochastic masking of biochemical properties exposes the model to varying levels of contextual information, enabling it to produce high-fidelity sequences under full biochemical input and more exploratory solutions when those cues are attenuated. In this formulation, full-information recovery reflects how well the model internalizes structure–biochemistry relationships, whereas masking modulates the effective information available to the model and thus the degree of permissible sequence variation. Importantly, the stochastic masking described here is applied uniformly and at random during training to build robustness across information regimes. In practical design workflows, users may instead choose to selectively mask specific regions, for example, active sites or binding interfaces, to direct localized exploration.

Moreover, we demonstrate that modeling continuous biochemical properties improves diagnostic inverse folding performance and yields consistent gains across enzyme, peptide, and antibody design tasks, highlighting the practical value of biochemistry-aware structure representations.

## 2 Methodology

### 2.1 Input Representation

A protein is a three-dimensional macromolecule containing one or multiple polypeptide chains, each chain formed by amino acid residues. Given a protein structure, we represent it as a graph 𝒢 (𝒱, ℰ, ℱ_𝒱_, ℱ_ℰ_) according to the featurizer in UniIF [27]. In protein structures, biochemical properties can be viewed as spatially varying physicochemical quantities—such as local charge or hydrophobicity—defined throughout the three-dimensional protein volume. Formally, these quantities can be represented by a continuous function *ϕ* : ℝ^3^ → ℝ^*d*^, which assigns to each spatial location **x** ∈ ℝ^3^ a vector of physicochemical values. Thus, *ϕ*(**x**) denotes the value of the biochemical properties (e.g., charge) at location **x**, not the coordinate itself. To obtain a computationally tractable approximation of this underlying field, we discretize *ϕ* via point sampling and feature assignment, capturing both surface and interior physicochemical patterns essential for protein sequence design.

Among various biochemical properties, we specifically choose hydrophobicity and charge to construct our biochemical features, according to SurfPro [28], which demonstrated these two properties to be the most informative for protein sequence design. Our feature construction process consists of four key stages:

**(i) Surface Representation:** We first generate a simplified-protein surface representation using MSMS (based just on alpha carbon spheres), which constructs a triangulated mesh based on the solvent-accessible surface area (SASA) model. The vertices of this mesh form an initial surface point cloud 𝒫_*S*_, which undergoes Gaussian smoothing followed by octree-based compression to ensure high-quality and uniform point distribution along the protein surface. (ii) **Internal Space Sampling:** To capture the biochemical environment within the protein core, we construct a bounding box encompassing the protein structure and uniformly sample 5000 points within this volume. We retain only those points that fall within the protein interior, as determined by a Delaunay triangulation (and associated Voronoi construction) of 𝒫_*S*_, forming the internal point cloud 𝒫_*I*_. (iii) **Point Cloud Integration:** The surface (𝒫_*S*_) and internal (𝒫_*I*_) point clouds are merged and subsampled to create a unified representation of 5000 points that captures both surface and internal biochemical environments. (iv) **Biochemical Feature Assignment:** Once the unified point cloud is constructed, we need to associate relevant biochemical properties with each point to create a meaningful representation of the protein’s chemical environment. For each point in the unified cloud *P*_*i*_ ∈ 𝒫, we determine its nearest residue by computing distances to all C_*α*_ atoms in the protein structure. Specifically, we identify the residue *r*_*j*_ whose C_*α*_ position **c**_*j*_ minimizes the Euclidean distance ∥ **x**_*i*_ − **c**_*j*_ ∥_2_ to point **x**_*i*_. We then transfer the biochemical properties of this nearest residue to the point. Each point is decorated with two key biochemical features: hydrophobicity *h*(**x**_*i*_) and charge *c*(**x**_*i*_) values derived from standardized amino acid property tables (detailed in Appendix A). For example, if a point is nearest to an isoleucine residue, it would be assigned a high hydrophobicity value of 4.5 and a neutral charge value of 0, whereas a point nearest to an arginine residue would receive a highly hydrophilic value of −4.5 and a positive charge value of +1. This assignment process creates a continuous representation of biochemical properties across both the protein surface and interior, where points in spatial proximity to similar residues will exhibit similar property distributions. The resulting attributed point cloud 𝒫 = {*P*_1_, *P*_2_, …, *P*_5000_ } consists of points *P*_*i*_ = {**x**_*i*_, *ϕ*(**x**_*i*_) }, where **x**_*i*_ ∈ ℝ^3^ represents spatial coordinates and *ϕ*(**x**_*i*_) = (*h*(**x**_*i*_), *c*(**x**_*i*_)) encodes the hydrophobicity and charge values. This approach transforms discrete residue-level biochemical properties into a continuous field distributed throughout the protein’s three-dimensional structure, providing a more natural representation of the biochemical environment as it exists in the folded protein state.

### 2.2 Model Architecture

#### 2.2.1 Struct-Encoder

To effectively capture both local and global structural information, we augment the protein structure graph with auxiliary nodes that serve as dedicated feature aggregators at different spatial scales. Specifically, we introduce two types of aggregator nodes: (1) a global aggregator that summarizes features of the entire protein structure and (2) several local aggregators that capture structural patterns within specific spatial regions. This hierarchical aggregation scheme is inspired by the [CLS] (classification) token mechanism in Transformer models, where special tokens accumulate and summarize information from input sequences.

Formally, we augment the original structure graph 𝒢 into an enhanced graph 𝒢^′^(𝒱^′^, ℰ^′^, ℱ_𝒱_, ℱ_ℰ_) by adding special aggregator nodes. The augmented node set is defined as 𝒱^′^ = 𝒱∪ 𝒱_*l*_ ∪ {*µ*}, where 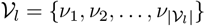 represents a set of local aggregator nodes that summarize structural information from specific regions of the protein, and *µ* is a global aggregator node that captures protein-wide features. In our work, we use |𝒱_*l*_ | = 8 local aggregator nodes.

To enable effective feature aggregation, we construct additional edges ℰ_*c*_ to connect these aggregator nodes to the protein structure: we randomly select |𝒱_*l*_ | centers 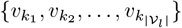 from the original nodes, and each local aggregator *ν*_*i*_ is connected to all nodes within a sphere region ℛ_*i*_ of radius *r*_*L*_ centered at its corresponding 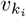. This creates local receptive fields where each aggregator *ν*_*i*_ collects and summarizes information from a specific spatial region of the protein. Meanwhile, the global aggregator *µ* is connected to all nodes in 𝒱 to capture protein-level features without spatial restrictions. The complete edge set is thus ℰ^′^= ℰ ∪ ℰ_*c*_.

After constructing 𝒢, UniIF is adapted to process it into node embeddings.

#### 2.2.2 The BC-Encoder

Similar to the introduction of aggregator nodes in Sec. 2.2.1, the local biochemical aggregator points, 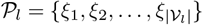, and a global biochemical aggregator point, *η*, are added to the point cloud 𝒫 together for each protein. Specifically, we define a local biochemical aggregator point *ξ*_*i*_ in the Struct-Encoder for each spherical region ℛ_*i*_ ⊂ ℝ^3^. Denote the augmented point cloud as 𝒫^′^= 𝒫 ∪ 𝒫_*l*_ ∪ {*η*}. Similar to the structure aggregator nodes, the embeddings of the biochemical aggregator points are initialized as learnable parameters 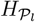 and *H*_*η*_. For each point *P*_*i*_ ∈ 𝒫, their features *ϕ*(**x**_*i*_) are passed through an FFN, resulting in a set of embeddings *H*_𝒫_. We then define the augmented embeddings as 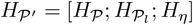.

The encoder processes the augmented point cloud using a multi-scale approach (Fig. 2(b)). For each point *P*_*i*_ ∈ 𝒫, we define its multi-scale neighborhood, 𝒩_*r*_(*P*_*i*_), which consists of all points within a radius *r* and the aggregator nodes *ξ*_*k*_ and *η*: 𝒩_*r*_(*P*_*i*_) = { *P*_*j*_ | ∥**x**_*i*_ − **x**_*j*_∥ ≤ *r*} ∪ { *ξ*_*k*_ | *P*_*i*_ ∈ ℛ_*k*_} ∪ {*η*}. This neighborhood defines the local context around point *P*_*i*_ at different spatial scales, where each radius *r* captures a different level of detail. Meanwhile, for each local aggregator nodes *ξ*_*k*_ and global node *η*, their neighborhoods are defined to contain themselves only: 𝒩_*r*_(*ξ*_*k*_) = {*ξ*_*k*_} and 𝒩_*r*_(*η*) = {*η*}.

Next, feature aggregation is performed using multi-head attention (MHA) applied to the embeddings of the point 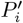 and its neighborhood 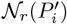. The result of the MHA operation is pooled using mean pooling to obtain the aggregated feature for each scale: 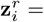 MeanPool(MHA 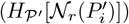). This operation is repeated for multiple radii, creating a set of aggregated features, each corresponding to a different scale. To combine these multi-scale features, we concatenate the aggregated features from different radii: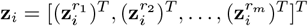. In our work, the number of scales, *m*, is set to 4. Finally, the fused feature **z**_*i*_ is passed through an FFN surrounded by a residual connection and followed by a linear transformation, yielding the final output: 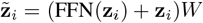.

**Fig. 2:**
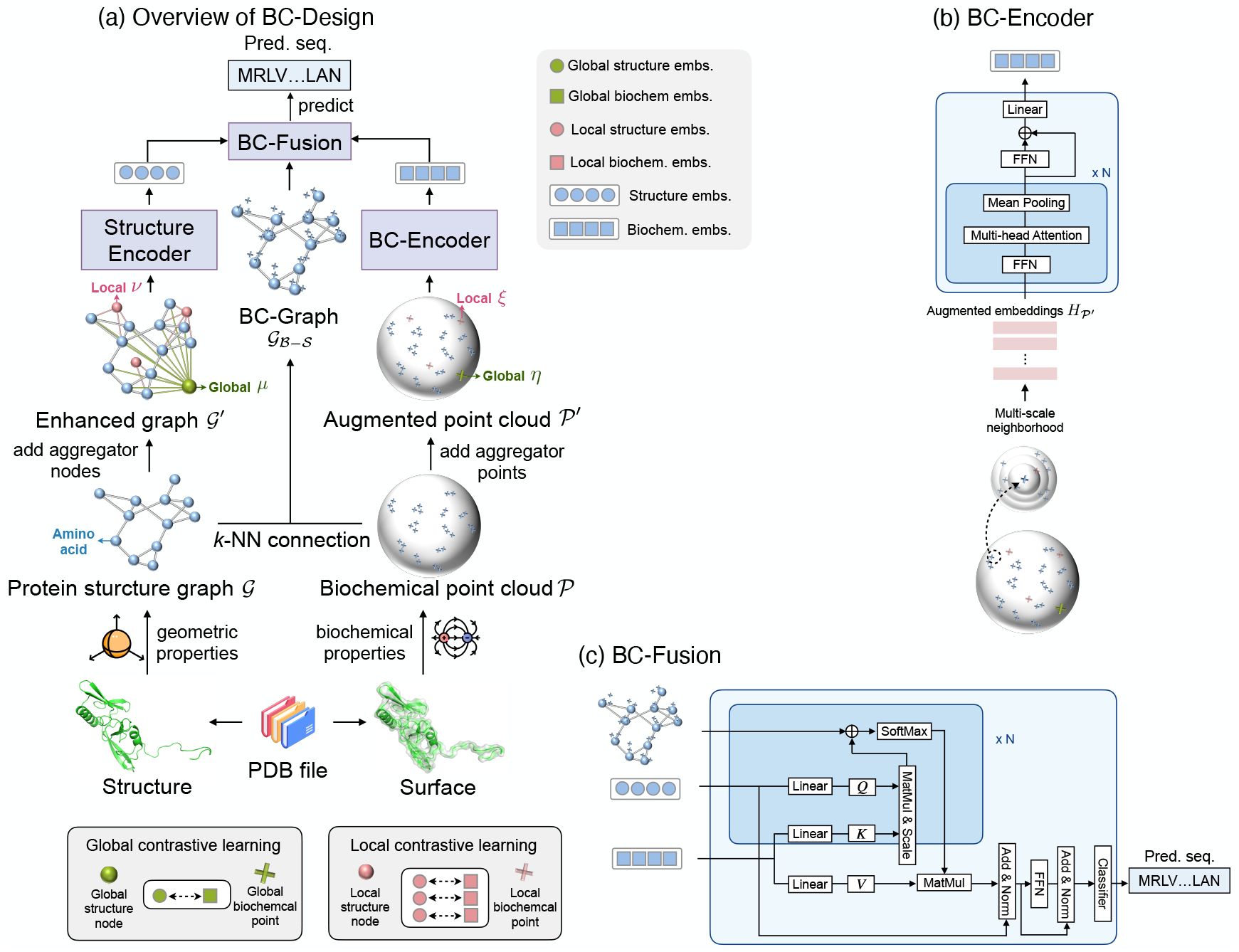
Overview of our proposed BC-Design framework. **(a)** Architecture and workflow: The framework takes protein backbone structures (PDB files) as input and predicts amino acid sequences (MRLV…LAN) as output. The structure is represented as a graph 𝒢 and enhanced with aggregator nodes (green: global structure *µ*, pink: local structure *ν*) to form 𝒢^′^. Similarly, biochemical features (hydrophobicity and charge) are represented as a point cloud 𝒫 augmented with aggregator points (green: global biochemical *η*, pink: local biochemical *ξ*) to form 𝒫^′^. The Struct-Encoder processes 𝒢^′^ to produce structure embeddings (blue circles), while the BC-Encoder processes ^′^ to generate biochemical embeddings (blue squares). These embeddings are integrated through BC-Fusion using the bipartite BC-Graph (𝒢_*B*−*S*_) to predict sequences. During training, contrastive learning aligns global structure embeddings with global biochemical embeddings (left box), and local structure embeddings with local biochemical embeddings (right box), enabling the model to function with only structural input during inference. **(b)** The BC-Encoder employs multi-scale neighborhood sampling and multi-head attention to capture biochemical property distributions, producing augmented embeddings 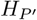. **(c)** BC-Fusion combines structure and biochemical embeddings through masked Transformer decoder layers guided by the BC-Graph, ultimately predicting the amino acid sequence through feedforward networks and a softmax layer.

#### 2.2.3 BC-FusionDecoder and Residue Classification

BC-Fusion serves as a specially designed decoder to integrate structural and biochemical information for amino acid sequence prediction in three steps (Fig. 2(c)):

First, it establishes spatial correspondences between structural and biochemical features through a bipartite graph structure (BC-Graph). Specifically, the BC-Graph 𝒢_ℬ−𝒮_ (𝒱_ℬ_,ℰ_ℬ_) connects each residue node in 𝒱 to its *k*_ℬ_ nearest points in the biochemical point cloud 𝒫, where 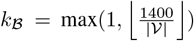 adapts to protein size. Second, it performs feature fusion using masked transformer decoder layers, where the attention mechanism is guided by the BC-Graph’s adjacency matrix. This ensures that each residue position attends only to its relevant biochemical features during decoding. Finally, for each residue position, the fused features are transformed through an FFN and softmax layer to predict probabilities over the 20 standard amino acids: 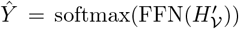, where 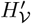 represents the fused embeddings at each residue position. This hierarchical decoding process enables the model to leverage both structural context and spatially aligned biochemical properties when predicting amino acid identities.

### 2.3 Training Strategy

We adopt a staged training strategy that progressively couples the structural and biochemical components of the model (Fig. 2(a)). We begin by pretraining the Struct-Encoder alone on the sequence prediction objective to establish a strong geometric backbone. Once this module is stabilized, its parameters are frozen and the BC-Eencoder is trained separately using the global and local contrastive objectives described below. After both encoders have been independently optimized, we freeze them and train the BC-Fusion decoder and prediction head to integrate the two modalities for sequence prediction. Finally, the full model is fine-tuned end-to-end on the same task, enabling joint optimization of structure, biochemical, and fused representations.

#### 2.3.1 Loss Function

##### Primary Sequence Prediction Objective (training stage (1), (3), and (4))

The sequence prediction objective is formulated as a cross-entropy loss: 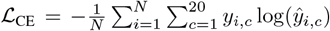, where *N* is the number of residues, *y*_*i,c*_ is the ground truth one-hot encoding for residue *i* and amino acid class *c*, and *ŷ*_*i,c*_ is the predicted probability for residue *i* being amino acid *c*.

##### Global Contrastive Learning (GCL)

For each protein structure, we form positive pairs using its global structure aggregator node and global biochemical aggregator point. To construct negative pairs, we maintain two continuously updated queues, 𝒬_𝒮_ and 𝒬_ℬ_, which store global structure embeddings and global biochemical embeddings, respectively, from the last *K*_ℬ_ = 64 proteins encountered during training. The global contrastive loss is defined as:

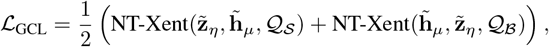

where NT-Xent is the normalized temperature-scaled cross-entropy loss used in contrastive representation learning. For an anchor embedding *u*, a positive embedding *v*, and a set of negative embeddings {*n*_*j*_}, NT-Xent is defined as:

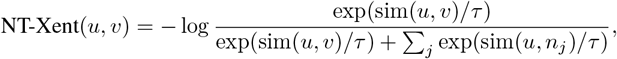

where sim(·, ·) denotes cosine similarity and *τ* is a temperature hyperparameter. This objective pulls the anchor *u* toward its positive partner *v* in embedding space while pushing it away from all negatives *n*_*j*_. In our setting, the positive pair consists of the global structure embedding 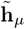 and the global biochemical embedding 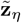 of the same protein, and the negative samples are embeddings of other proteins stored in the queues. Thus, the contrastive objective drives corresponding structure–biochemistry representations to align while ensuring that embeddings from different proteins remain well separated.

##### Local Contrastive Learning (LCL)

At the local level, the non-corresponding node-point pairs within the same protein are treated as negative pairs. Since biochemical properties like hydrophobicity and charge are inherent to amino acids and less variable than structural conformations, we encourage the local structure embeddings to learn and reflect essential biochemical information. To adjust the local structure embeddings to better align with the corresponding local biochemical embeddings and preserve the local biochemical embeddings as stable representations of biochemical properties, we design asymmetric contrastive learning by only using local structure embeddings as anchors in the NT-Xent losses. 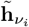 and 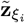 are the local structure and local biochemical embeddings, respectively, the local contrastive loss is defined as:

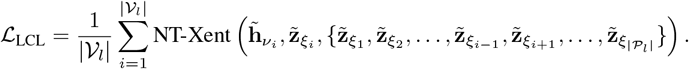

##### Combined Loss Function (training stage (2))

In stage 2, the model is trained using a combined loss function: ℒ = *λ*_1_ ℒ_GCL_ + *λ*_2_ ℒ_LCL_, where *λ*_1_ = 0.01 and *λ*_2_ = 1 are the weights for the global and local contrastive losses. This combined objective ensures that the model maintains consistency between structural and biochemical representations at both global and local scales before stage 3 and 4.

#### 2.3.2 Masking Strategy

During training, we apply stochastic masking independently to structural and biochemical inputs in order to promote robustness under missing-information regimes. Structural features are randomly masked with probability 0.1 whenever the Struct-Encoder is trainable. Biochemical features follow a mixed masking schedule during end-to-end training: with probability 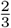, all biochemical inputs are fully masked (*p*_*b*_ = 1), while in the remaining cases the masking ratio is sampled from Uniform(0, 1). This strategy ensures robustness across varying information regimes. While full biochemical properties yield the highest accuracy, the model remains functional under partial or absent biochemical inputs, an advantageous property for real-world scenarios.

## 3 Results

### 3.1 Main Performance Evaluation (CATH 4.2)

To assess the effectiveness of BC-Design in generating protein sequences from backbone structures and biochemical feature distributions, we evaluate its performance against recent state-of-the-art methods on the **CATH 4.2** dataset [29]. This comprehensive dataset encompasses complete protein chains up to 500 residues in length, with structures organized at 40% non-redundancy based on their CATH classification (Class, Architecture, Topology, Homologous). Following the protocol established in previous work [18], we adopt an identical data split comprising **18**,**024** training, **608** validation, and **1**,**120** test proteins, with no CAT overlap across sets.

To comprehensively evaluate BC-Design’s performance, we conduct both sequence-level and structure-level validations. At the sequence level, we benchmark against a wide spectrum of state-of-the-art methods, including traditional graph-based approaches (GVP [30], StructGNN [30], GraphTrans [18], GCA [31]), recent advanced architectures (ProteinMPNN [19], PiFold [2], AlphaDesign [32], SPIN-CGNN [33], GRADE-IF [34], DIPRoT [35], SPDesign [36], ProRefiner+ESM-IF1 [37]), and language model-based methods (ESM-IF1 [4], LM-Design [38]). We also compare with methods incorporating external knowledge or multi-modal learning (Knowledge-Design [39], MMDesign [40], VFN-IFE [41]). For structure-level assessment, we evaluate against leading approaches, including ESM-Design [42], AlphaFold-Design (AF-Design) [43], PiFold, GraphTrans, GVP, ByProt [44], and ProteinMPNN.

#### Sequence-level validation

To evaluate BC-Design’s performance in protein design, we employ three complementary metrics on the CATH 4.2 test set [45]: sequence recovery, perplexity, and native sequence similarity recovery (nssr) [46]. Sequence recovery quantifies the model’s accuracy in reproducing native amino acid sequences by calculating the percentage of positions where the predicted amino acid matches the native one. Perplexity, defined as 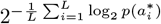 where 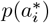 is the predicted probability of the native amino acid 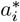 at position *i* in a sequence of length *L*, measures the model’s uncertainty in its predictions. Intuitively, perplexity represents the effective number of amino acids the model considers at each position—a perplexity of 2.0 indicates that the model is as uncertain as if it were randomly choosing between two equally likely amino acids at each position. The ideal perplexity is 1.0 (perfect certainty), while higher values (up to 20 for completely random predictions across all amino acids) indicate increasing uncertainty. Perplexity is dimensionless and serves as a more sensitive measure of prediction quality than sequence recovery alone, as it accounts for the model’s confidence in its predictions even when they don’t exactly match the native sequence. The nssr metric extends beyond exact matches by considering residue similarity based on BLOSUM62 substitution scores [47], counting positions as correctly predicted if the model generates an amino acid that is biochemically similar to the native one, thereby providing a more functionally relevant evaluation of sequence design quality.

Although recovery is a standard diagnostic metric in inverse folding, it should not be interpreted as a complete measure of design quality. Many positions in natural proteins admit multiple biochemically plausible residues, meaning that matching the native residue often reflects correct modeling of structural and physicochemical constraints rather than memorization of the wild-type sequence. Consequently, recovery primarily captures how well a model internalizes structure–biochemistry relationships, not whether it reproduces the exact native sequence. To obtain a more comprehensive assessment of design performance, we therefore complement recovery with perplexity, nssr, and structure-based evaluations.

We benchmark BC-Design against leading approaches, including ProRefiner+ESM-IF1, ESM-IF1, GVP, ProteinMPNN, and ProteinMPNN-C, specifically for the nssr metric. As shown in Fig. 3 (a, b), BC-Design significantly outperforms existing methods, achieving **90.25%** sequence recovery, **1.38** perplexity, and **94.71%** nssr score. These results demonstrate that our integrated approach of modeling both backbone structure and biochemical features substantially enhances protein design accuracy across diverse proteins in the CATH 4.2 dataset.

**Fig. 3:**
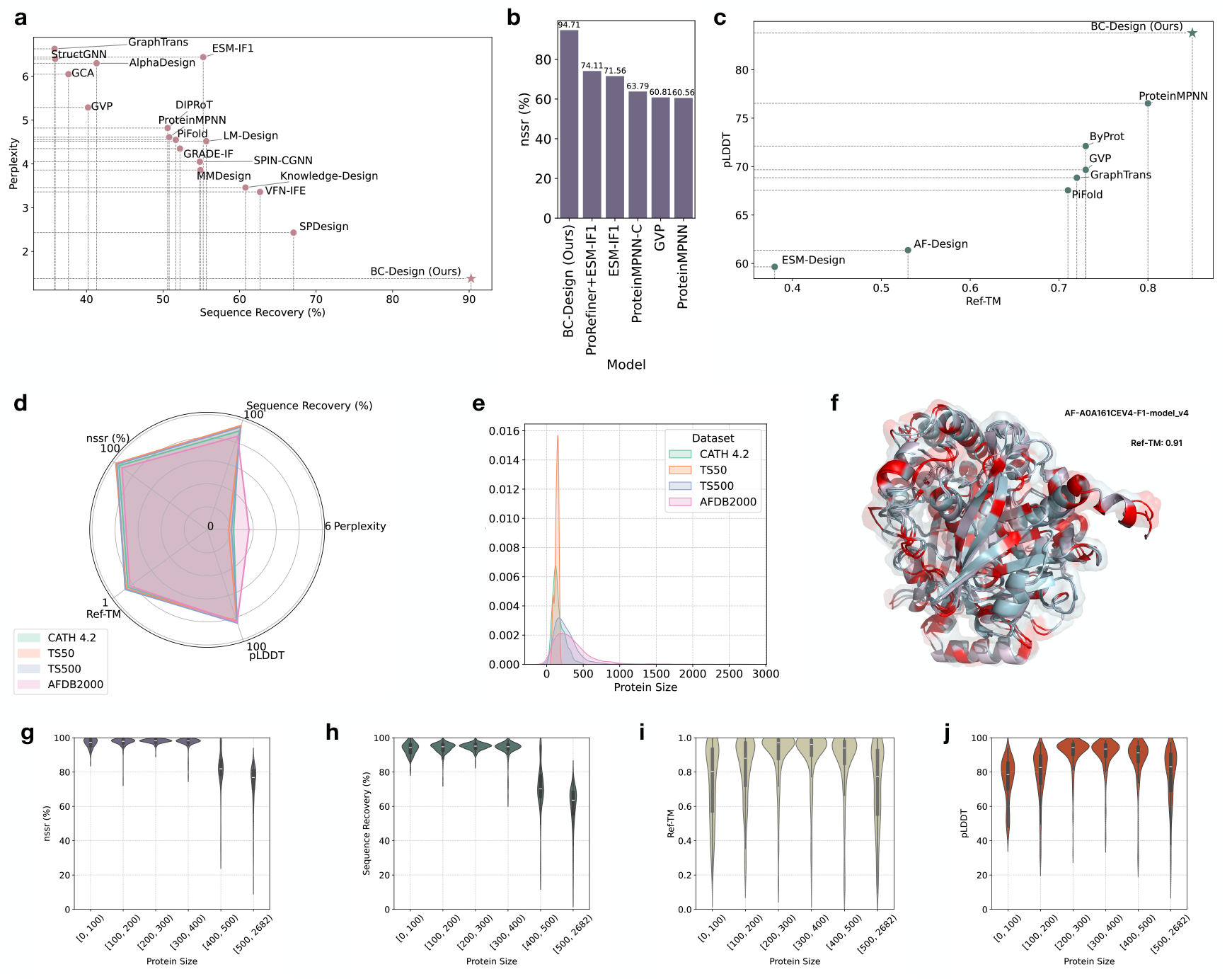
**(a)** Sequence recovery and perplexity on the CATH 4.2 test set. BC-Design achieves 90% recovery and the lowest perplexity among all models, illustrating a strong inverse correlation between these metrics and highlighting its superior sequence prediction capability. **(b)** Nssr comparison on CATH 4.2. BC-Design attains 94.71% nssr, markedly outperforming ProteinMPNN-C, ESM-IF1, and ProRefiner+ESM-IF1, indicating more biochemically consistent substitutions and improved functional plausibility. **(c)** Structural quality of ESMFold-refined predictions on the filtered CATH 4.2 set. BC-Design preserves high Ref-TM and pLDDT scores, whereas models such as ESM-Design and AF-Design show substantial structural degradation. **(d)** Cross-dataset performance. BC-Design generalizes well across CATH 4.2, TS50, and TS500; performance on AFDB2000 shows moderate sequence decline but strong structural fidelity. **(e)** Protein length distributions across datasets. AFDB2000 contains substantially more long proteins (*>*500 aa), explaining the distributional shift observed in (d). **(f)** Case study of AF-A0A161CEV4-F1-model v4 (622 aa). BC-Design achieves a Ref-TM of 0.91 and accurately recovers hydrophobicity patterns despite the sequence length exceeding typical training ranges. **(g–j)** Performance stratified by protein size on AFDB2000. Sequence recovery decreases for proteins *>*400 aa, while structural metrics peak at medium lengths and remain reasonably high even for large proteins, indicating that BC-Design maintains fold-level accuracy under substantial distributional shift.

#### Structure-level validation

While sequence-level metrics provide valuable insights, they cannot fully capture the practical effectiveness of protein design models. Given that protein function critically depends on structure, we perform a structural evaluation of our designs. We utilize ESMFold [48] for structure prediction due to its comparable accuracy to AlphaFold 2 [8] with improved computational efficiency. Following [49], we use a curated subset of **82** proteins from the CATH 4.2 test set by selecting one protein from each CATH family randomly. The structural quality is assessed using two complementary metrics: Ref-TM [49] for structural similarity to native conformations, and pLDDT [8] for prediction confidence. As shown in Fig. 3 (c), BC-Design achieves superior structural accuracy compared to baseline methods, with Ref-TM of **0.85** and pLDDT of **83.79**. pLDDT (predicted Local Distance Difference Test) is a confidence score introduced in AlphaFold2 that measures the expected accuracy of predicted structural elements on a per-residue basis, ranging from 0 to 100. Higher pLDDT values indicate greater confidence in the structural prediction—scores above 70 generally correspond to high-confidence regions with reliable atomic positions, while values below 50 indicate regions of low confidence. The high average pLDDT of 83.79 achieved by our method suggests that the proteins designed by BC-Design fold into well-defined structures with high confidence, indicating not just sequence recovery but functional structural formation. These results suggest that BC-Design effectively captures the complex relationships between sequences, structures, and biochemical features, rather than merely optimizing sequence similarity. The reliability of these results is reinforced by [49], which demonstrates consistent performance rankings across different structure prediction models, including ESMFold, AlphaFold 2, and OmegaFold [50].

To contextualize these structure-level metrics, we also quantify the empirical upper bound imposed by the structure predictor itself. Specifically, we fold the native wild-type sequences from the CATH 4.2 test set using ESMFold and compare the resulting structures to their corresponding crystal structures. Under this evaluation protocol, native sequences achieve an average pLDDT of approximately 85 and a Ref-TM of 0.86 (Table 1). These values represent the highest scores achievable with ESMFold, as even the correct native sequences do not yield perfect agreement with crystal structures due to inherent limitations in structure prediction.

**Table 1:** Upper bounds for structural metrics established by folding native wild-type sequences with ESMFold.

| Dataset | pLDDT | Ref-TM |
| --- | --- | --- |
| CATH 4.2 test set | 85 | 0.86 |
| AFDB2000 | 85 | 0.83 |
| TS50 | 83 | 0.87 |
| TS500 | 86 | 0.89 |

The structural accuracy attained by BC-Design (pLDDT 83.79; Ref-TM 0.85) therefore lies very close to this empirical ceiling. Importantly, in inverse folding the ground-truth structure is the native backbone itself; the goal is to assess whether a designed sequence folds back into this target structure family. Accordingly, we fold each designed sequence using ESMFold and compare the predicted structure to the native backbone, following standard practice in inverse design. The close match between BC-Design’s folded structures and the native backbone indicates that its designed sequences achieve nearly the same level of structural confidence and fidelity that modern predictors obtain on the true sequence. This supports the conclusion that the model is learning meaningful sequence–structure– biochemistry relationships rather than relying on shortcut signals or trivially recovering native residues.

### 3.2 Why Does It Work?

#### Input ablation: roles of structure, biochemistry, and fusion

We conducted an ablation study to assess the impact of different input components on sequence recovery, as summarized in Fig. 4(c). Four main observations emerge from this analysis. First, each input stream makes a substantial contribution to overall performance; removing any component leads to a clear drop in sequence recovery. Second, biochemical features exert the strongest effect, yielding an improvement of approximately **41** percentage points in sequence recovery relative to models that rely solely on geometric information. Third, removing the BC-Graph fusion module and replacing it with a generic Transformer decoder results in a similar **41** percentage point decrease in recovery, even though the underlying inputs remain essentially unchanged. This highlights that dedicated cross-modal fusion, rather than simply concatenating or jointly encoding inputs, is crucial for extracting value from the biochemical properties. Fourth, the additional gain from including backbone structural features is more modest (around **9** percentage points) but still meaningful, indicating that geometry alone cannot explain the model’s performance, yet it provides complementary signals to the biochemical properties.

**Fig. 4:**
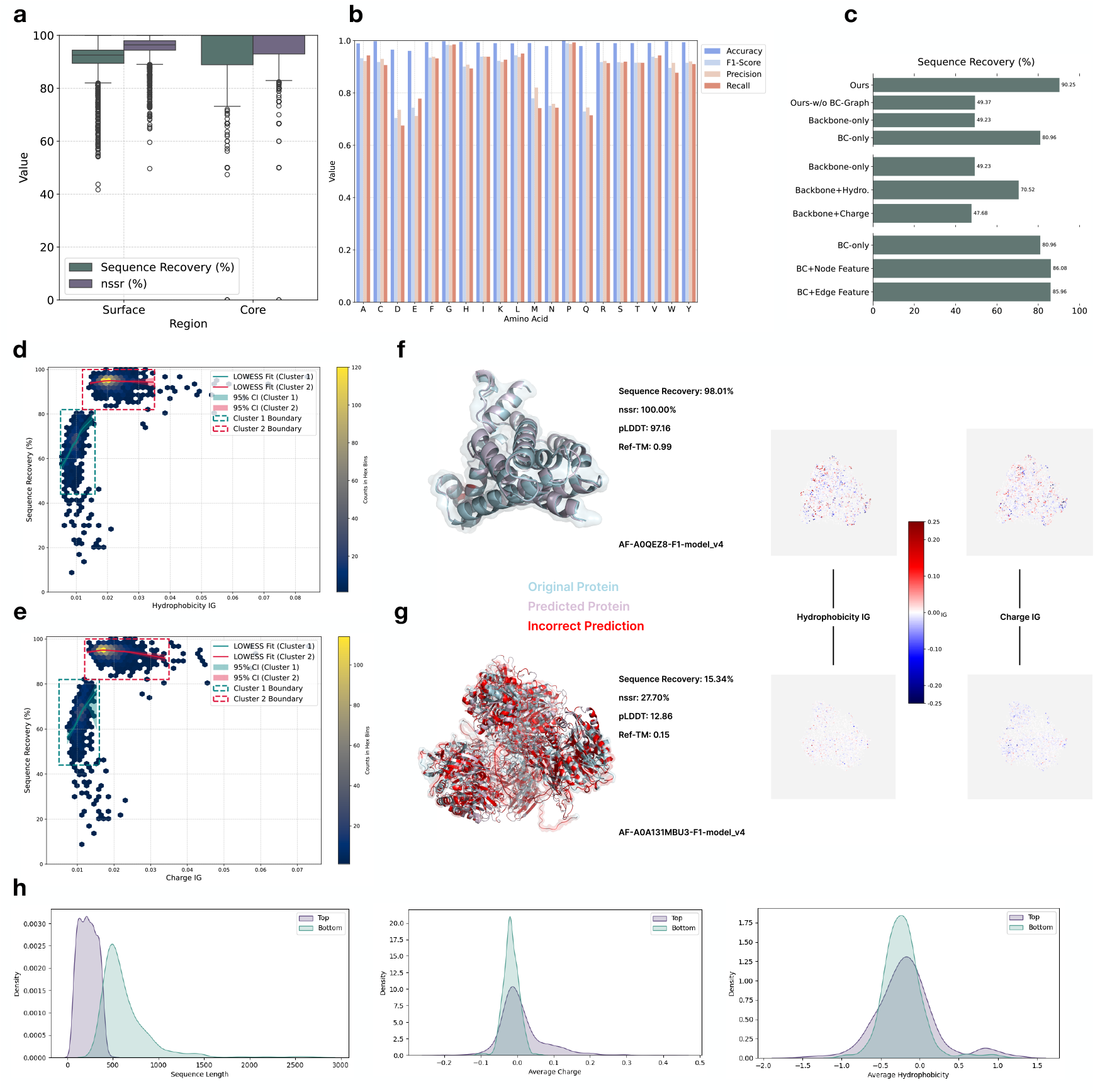
**(a)** Sequence recovery and nssr for surface vs. core residues. Core positions exhibit substantially higher accuracy, reflecting stronger structural constraints in protein interiors. **(b)** Residue-level precision and recall across amino acid types. Negatively charged (D, E) and polar amides (N, Q) show lower performance due to biochemical similarity and frequent localization in flexible surface regions. **(c)** Ablation results quantifying contributions of each component. Removing biochemical features leads to the largest drop in accuracy, followed by replacing the BC-Graph module, while structural features provide a smaller but meaningful gain. **(d**,**e)** Relationship between sequence recovery and IG attribution for hydrophobicity and charge. Two clusters emerge: low-recovery proteins with consistently low IG values, and high-recovery proteins with higher attribution. LOWESS fits show strong dependence on biochemical cues in the low-recovery regime and diminishing returns at higher IG. **(f**,**g)** Case studies illustrating attribution patterns in successful vs. failed predictions. High-recovery, high–Ref-TM examples show strong biochemical feature attribution, whereas poorly predicted proteins exhibit uniformly weak IG signals. **(h)** Distributions of sequence length, average charge, and hydrophobicity for the two clusters in (d,e), highlighting that low-recovery cases tend to be longer and biochemically more homogeneous.

#### Masking-based quantification of biochemical dependence

To further quantify the role of biochemical properties, we performed an inference-time masking experiment in which we progressively withheld a fraction of the biochemical point cloud. This procedure directly modulates the amount of biochemical information available to the model and allows us to probe how sequence recovery, foldability, and sequence diversity respond to systematic removal of biochemical cues.

#### Motivation

This experiment is designed to characterize the dependence of BC-Design on biochemical information from a functional standpoint. By gradually masking the hydrophobicity and charge properties, we can trace how the model transitions between high-fidelity reconstruction and exploratory design regimes, and how these changes affect structural quality.

#### Method

Masking ratios ranged from 0% (full biochemical information) to 100% (no biochemical information). For a given masking ratio, a corresponding fraction of points in the biochemical point cloud was selected at random, and their true feature values were replaced with a learnable mask token. All experiments were conducted on the CATH 4.2 test set under the same decoding protocol used in the main evaluation.

#### Metrics

Recovery and Ref-TM were computed from greedy decoding for each protein. Sequence diversity was estimated by sampling *N* = 20 sequences per protein at temperature 1.0 and averaging the pairwise Hamming distance across sampled sequences:

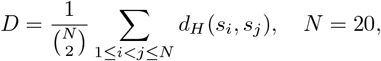

where *d*_*H*_ denotes the Hamming distance between sequences *s*_*i*_ and *s*_*j*_.

As summarized in Table 2, sequence recovery decreases smoothly and predictably as the masking ratio increases, confirming that biochemical properties are essential for achieving high accuracy. At the same time, sequence diversity increases monotonically, while structural quality, as measured by Ref-TM, remains high and only gradually declines. This behavior reveals a controllable recovery–diversity trade-off: by adjusting the availability of biochemical information, BC-Design can be tuned to operate either in a high-fidelity regime, where native-like sequences are prioritized, or in a more exploratory regime, where diverse yet foldable sequences are generated.

**Table 2:** Trade-off between sequence recovery, structural quality, and diversity under different biochemical masking ratios on the CATH 4.2 test set. As more biochemical information is masked, recovery decreases and diversity increases, while Ref-TM remains high, indicating that designed sequences continue to fold into native-like structures.

| Metric | Percentage of Masked Biochemical Feature |  |  |  |  |  |  |  |
| --- | --- | --- | --- | --- | --- | --- | --- | --- |
|  | 0% | 20% | 40% | 60% | 80% | 90% | 95% | 100% |
| Sequence Recovery (%) | 90.25 | 89.91 | 88.94 | 86.57 | 79.83 | 70.74 | 62.31 | 49.23 |
| Ref-TM | 0.85 | 0.85 | 0.85 | 0.85 | 0.84 | 0.83 | 0.82 | 0.81 |
| Diversity (%) | 11.36 | 11.54 | 12.88 | 15.87 | 23.56 | 32.93 | 41.05 | 51.49 |

This tunability illustrates an additional functional benefit of biochemical augmentation. Beyond boosting raw accuracy, continuous biochemical properties give BC-Design a principled mechanism to navigate the design space: the same model can be used either to produce conservative, native-like sequences or to explore diverse alternatives that remain structurally plausible.

#### Attribution analysis with integrated gradients

BC-Design leverages both structural and biochemical information, and the ablations above indicate that biochemical properties have a dominant effect on sequence prediction. To better understand this observation, we applied integrated gradients (IG) [51] to quantify how hydrophobicity and charge influence the model’s predictions on proteins from AFDB2000.

Integrated gradients (IG) is an attribution method that assigns an importance score to each input feature by integrating gradients along a straight-line path from a baseline input to the actual input. Formally, for feature *i*,

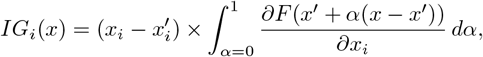

where *x* is the input, *x*^′^ is a baseline, and *F* (·) denotes the model output with respect to which attributions are computed. In our case, *F* corresponds to the model’s predicted logit for the residue type at the target position, ensuring that IG measures how each biochemical feature influences the model’s sequence prediction.

For each protein, we compute IG scores for hydrophobicity and charge at every point in the biochemical point cloud with respect to the predicted probabilities of the native amino acids. This yields two IG point clouds per protein, one for each biochemical feature. We then aggregate these scores by taking the average absolute IG value per feature, producing a global importance score for hydrophobicity and charge for each protein. Finally, we examine how these scores relate to sequence recovery, as shown in Fig. 4(d,e).

A clear pattern emerges. Proteins with high sequence recovery (for example, above 80%) almost always exhibit high IG scores for both hydrophobicity and charge. Conversely, proteins with low sequence recovery consistently display low IG values. This stratification indicates that successful sequence reconstruction is tightly coupled to the effective use of biochemical information: whenever BC-Design recovers sequences well, biochemical properties make strong, measurable contributions; when recovery is poor, the model fails to exploit these cues effectively. The strong correlation between IG magnitude and prediction success provides direct evidence that treating biochemical properties as continuous distribution on protein surfaces is central to the model’s performance.

Fig. 4(d,e) further reveals a distinctly bimodal structure in the IG–recovery landscape. We observe one cluster with low recovery and low IG (roughly below 0.016), and another with high recovery and higher IG values. This motivates two complementary threshold concepts: a recovery threshold effect and an IG threshold effect.

For the **recovery threshold effect**, we find that proteins with recovery below 80% almost exclusively occupy the low-IG regime. In this range, increases in IG are associated with steep gains in recovery, indicating that biochemical cues are the limiting signal and that better utilization of hydrophobicity and charge can substantially improve performance. In contrast, for proteins with recovery above 80%, IG values are generally higher but exhibit little correlation with further recovery improvements, suggesting a regime of diminishing returns once sufficient biochemical evidence has been captured.

For the **IG threshold effect**, IG values below approximately 0.013 are almost always associated with recovery below 80%, whereas IG values above roughly 0.016 correspond to consistently high recovery. Between these thresholds lies a transition region where recovery varies more widely, reflecting intermediate levels of biochemical influence. This pattern suggests that below a certain level of biochemical contribution the model is unable to resolve residue identities reliably, while beyond a higher threshold additional biochemical signal yields only marginal benefits.

To investigate the origin of these regimes, we conducted a cluster-level analysis of the IG–recovery landscape. The IG distributions show a striking bimodality (Fig. 4(d,e)), indicating that proteins fall into two biophysically distinct groups that differ in sequence length and global biochemical diversity.

#### Sequence length

As shown in Fig. 4(h), proteins in the high-recovery, high-IG cluster (Cluster 2, Top) are predominantly shorter than 500 residues. In contrast, proteins in the low-recovery, low-IG cluster (Cluster 1, Bottom) are substantially longer on average. This gives rise to two prediction regimes. Longer proteins reside in a <u>signal-dependent</u> regime, where increased structural complexity creates ambiguity that must be resolved by strong biochemical cues, leading to a steep IG–recovery relationship. Shorter proteins lie in a <u>signal-saturated</u> regime, where simpler geometry and clearer biochemical properties provide sufficient evidence for accurate predictions, rendering additional IG increases less impactful.

#### Biochemical diversity

Analysis of average hydrophobicity and charge (Fig. 4(h)) shows that proteins in the low-recovery cluster tend to be biochemically homogeneous, with global hydrophobicity and charge clustered near neutral values. In contrast, the high-recovery cluster exhibits greater biochemical diversity, with more pronounced global biases. When global biochemical signals are weak or ambiguous, the model lacks sufficient context to disambiguate residue identities, mirroring the low-IG, low-recovery regime.

These observations highlight that both hydrophobicity and charge play critical roles in determining prediction success and further support the conclusion that biochemical properties are central to BC-Design’s performance. They also imply that the model’s behavior is highly predictable from the strength of biochemical contributions: once IG scores fall below the identified thresholds, substantial drops in recovery are expected.

To illustrate these effects more concretely, we visualize two representative proteins, each shown with its native and predicted structures and corresponding IG distributions (Fig. 4(f,g)). In the well-predicted example (AF-A0QEZ8-F1-model v4), BC-Design achieves 98.01% sequence recovery and a Ref-TM of 0.99, accompanied by consistently large IG magnitudes across both biochemical features. In contrast, in the poorly predicted example (AF-A0A131MBU3-F1-model v4), recovery drops to 15.34% and Ref-TM to 0.15, and IG magnitudes are uniformly small. The stark difference between these cases reinforces that effective utilization of continuous biochemical properties is a key determinant of successful sequence prediction in BC-Design.

### 3.3 Robustness & Generalization

To evaluate the generalizability of BC-Design, we assessed its performance on three widely used test sets in addition to CATH 4.2: (i) **TS50**, a benchmark of 50 protein chains introduced in [14]; (ii) **TS500**, an expanded 500-protein benchmark derived in the same work; (iii) **AFDB2000**, a size- and function-agnostic collection of 2,000 proteins from the AlphaFold Database (AFDB) [52], selected alphabetically from Swiss-Prot to avoid biases induced by function or family. None of the AFDB2000 proteins overlap with the training or other evaluation sets used in this study.

#### Performance across heterogeneous datasets

As shown in Fig. 3(d), BC-Design performs consistently well on TS50 and TS500, achieving sequence- and structure-level metrics comparable to—or better than—those observed on the CATH 4.2 test set. On AFDB2000, however, sequence-level performance is reduced. To investigate this discrepancy, we examined the distribution of protein lengths across the four test sets (Fig. 3(e)) and found that AFDB2000 contains a substantially higher proportion of large proteins, many of which exceed the upper size range represented in the training data.

#### Understanding performance on AFDB2000

To quantify the effect of protein size, we partitioned AFDB2000 into six length bins: [0,100), [100,200), [200,300), [300,400), [400,500), and [500,+ ∞). As shown in Fig. 3(g,h), BC-Design maintains high sequence recovery and nssr for proteins shorter than 400 amino acids, matching performance on the other test sets. For larger proteins, sequence-level metrics decline, consistent with the limited representation of long proteins in the training distribution.

Structure-level metrics (Fig. 3(i,j)) reveal a different pattern. Ref-TM and pLDDT increase from small to medium sizes, peak around the [200,300) and [300,400) ranges, and then decline for longer proteins. Notably, the median Ref-TM values for the [0,100) and [400,500) bins are similar, indicating that even when sequence recovery decreases for large proteins, BC-Design still achieves reasonable folding accuracy. Several proteins in the largest bin (*>*500 residues) attain Ref-TM scores above 0.8. For example, for protein AF-A0A161CEV4-F1-model v4 (622 residues; Fig. 3(f)), BC-Design reaches a Ref-TM of 0.91, demonstrating robust structural generalization far beyond the training size regime.

These results indicate that the decline in sequence-level accuracy on AFDB2000 is driven by a distributional shift, specifically, the underrepresentation of large proteins in the training set, rather than a limitation of the architecture’s capacity to model long-range dependencies.

#### Augmenting training data to test architectural limits

To explicitly test whether the architecture can support large proteins when given appropriate training examples, we trained an enhanced model, BC-Design (LARGE), on an augmented dataset comprising the original CATH 4.2 training set plus 4,000 AFDB proteins between 400 and 1,000 residues. As summarized in Table 3, this strategy yields substantial improvements in the long-protein regime. For proteins longer than 500 residues, sequence recovery increases from 60.12% to 91.47%, and nssr shows a comparable gain. Structure-level metrics (Ref-TM and pLDDT) remain stable across both models, indicating that the underlying structural fidelity is preserved while sequence fidelity improves.

**Table 3:** Performance comparison on AFDB2000 between the original BC-Design and the augmented BC-Design(LARGE), showing substantial gains for proteins above 400 residues.

| Metric | Model | Sequence Length |  |  |  |  |  |
| --- | --- | --- | --- | --- | --- | --- | --- |
|  |  | [0, 100) | [100, 200) | [200, 300) | [300, 400) | [400, 500) | [500, +∞) |
| Sequence Recovery (%) | BC-DESIGN | 93.61 | 94.22 | 94.88 | 93.89 | 71.41 | 60.12 |
|  | BC-DESIGN (large) | 98.31 | 99.05 | 99.13 | 98.97 | 95.81 | 91.47 |
| nssr (%) | BC-DESIGN | 96.89 | 97.63 | 98.06 | 97.52 | 81.93 | 73.69 |
|  | BC-DESIGN (large) | 99.20 | 99.62 | 99.68 | 99.63 | 97.92 | 95.68 |
| Ref-TM | BC-DESIGN | 0.74 | 0.81 | 0.90 | 0.89 | 0.87 | 0.71 |
|  | BC-DESIGN (large) | 0.74 | 0.81 | 0.90 | 0.89 | 0.88 | 0.73 |
| pLDDT | BC-DESIGN | 75.43 | 78.89 | 91.38 | 90.47 | 87.37 | 78.37 |
|  | BC-DESIGN (large) | 75.21 | 78.83 | 91.41 | 90.49 | 89.73 | 81.90 |

Motivated by these observations, and to explicitly test whether the architecture itself can support large proteins when provided with appropriate training data, we trained an enhanced version, BC-Design (LARGE), on an augmented dataset. This expanded training set includes the original CATH 4.2 training set supplemented with 4,000 proteins between 400 and 1,000 residues from the AlphaFold Database. As shown in Table 3, this strategy substantially improves performance in the long-protein regime. In particular, sequence recovery for proteins longer than 500 residues increases from 60.12% to 91.47%, and nssr improves accordingly. These findings demonstrate that the architecture of BC-Design is fully capable of modeling large proteins and that performance in this regime can be readily extended through training on a more size-diverse dataset.

These findings demonstrate that BC-Design is architecturally capable of modeling proteins across a broad size spectrum and that limitations observed for large proteins arise primarily from the training distribution. Once supplied with more size-diverse data, the model generalizes effectively, achieving high sequence fidelity and maintaining strong structural accuracy.

#### Robustness to backbone coordinate noise

We further assessed robustness to structural imperfections by introducing controlled perturbations to backbone coordinates at inference time. Gaussian noise with standard deviations (*σ*) ranging from 0.02 Å to 1.0 Å was added independently to the N, C_*α*_, C, and O atoms, mimicking uncertainties commonly encountered in experimental structures and generative backbone models.

As shown in Table 4, BC-Design exhibits smooth, progressive degradation rather than abrupt failure. Performance remains largely intact under mild perturbations (*σ* = 0.02 Å, 88.00% recovery), and even under substantial noise (*σ* = 0.5 Å) the model retains close to 50% recovery. This behavior indicates that BC-Design does not rely on brittle geometric cues and can tolerate realistic levels of backbone variation.

**Table 4:** Sequence recovery on the CATH 4.2 test set under Gaussian perturbations applied to backbone atom coordinates. BC-Design exhibits smooth degradation and maintains substantial accuracy under realistic noise levels.

| Metric | Standard Deviation of Backbone Noise (Å) |  |  |  |  |  |  |  |  |
| --- | --- | --- | --- | --- | --- | --- | --- | --- | --- |
|  | 0 | 0.02 | 0.1 | 0.2 | 0.3 | 0.4 | 0.5 | 0.6 | 1.0 |
| Sequence Recovery (%) | 90.25 | 88.00 | 78.51 | 53.34 | 52.21 | 50.81 | 49.26 | 47.28 | 39.83 |

These results highlight an important aspect of the framework: continuous biochemical properties provide a stabilizing complementary signal that mitigates the impact of noisy or imperfect structural inputs. This robustness is particularly relevant for practical design workflows, where the target backbone is often not a high-resolution crystal structure but instead originates from low-resolution experiments, generative structure models, or molecular-dynamics snapshots with substantial conformational variability. In such settings, geometric information alone is typically insufficient, whereas biochemical properties offer a smooth, physically meaningful constraint that remains reliable even when the backbone is imprecise. As a result, BC-Design maintains stable and high-quality design behavior under realistic structural imperfections that arise routinely in forward protein engineering scenarios.

### 3.4 Practical Functionality in Enzyme, Peptide, and Antibody Design

We next asked whether the biochemistry-aware representations learned by BC-Design translate into practical gains in realistic protein engineering settings. To this end, we evaluated the model on three representative functional design tasks—enzyme redesign, peptide redesign, and antibody CDRH3 redesign—which probe distinct aspects of sequence–structure–function relationships beyond diagnostic inverse-folding metrics.

#### Enzyme substrate-binding design

To assess functional improvements in enzymatic activity, we applied BC-Design to redesign enzymes and quantified substrate-binding affinity using ESP scores [53]. The ESP (electrostatic potential) score is a functional metric that measures electrostatic complementarity between an enzyme and its substrate by comparing their surface potential fields; higher scores indicate stronger predicted binding affinity and improved catalytic compatibility. Across all tested substrates, BC-Design achieved the highest average ESP score (**0.9183**), outperforming ProteinMPNN [19], PiFold [2], LM-DESIGN [38], and SurfPro [54] (Table 5). The model also demonstrated strong zero-shot generalization, reaching an ESP score of **0.9841** on the unseen substrate C00001. Notably, even the backbone-only variant (BC-Design-BS), which does not access biochemical properties at inference, exceeded baselines on challenging substrates such as C00019. These results indicate that integrating continuous biochemical properties with geometric structure not only boosts diagnostic design metrics but also improves predicted catalytic competence across diverse substrate classes.

**Table 5:**
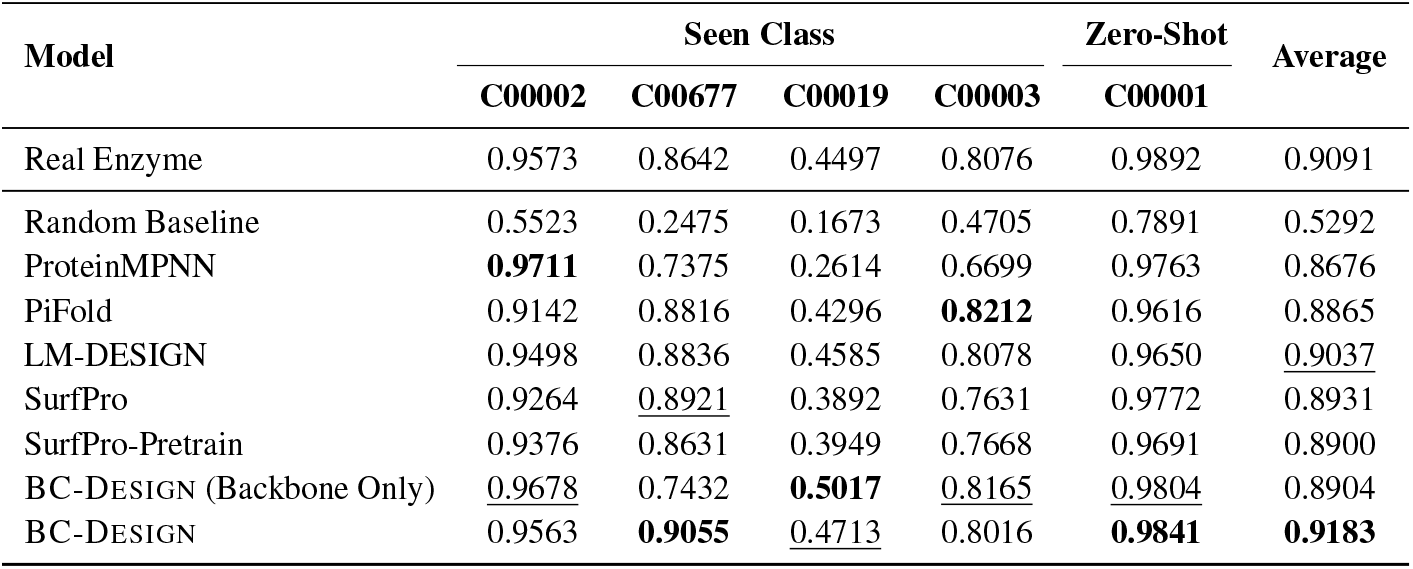
ESP scores (↑) of all models on the enzyme design task. Substrates are identified by KEGG IDs. “Average” denotes the ESP score over the full test set.

#### Peptide design

We next examined whether BC-Design can support peptide redesign under structural and functional constraints. On the PepMerge benchmark [55], using the preprocessing and sampling protocol from PepFlow, BC-Design outperformed all competing methods across sequence recovery, worst-case recovery, naturalness (Likeness), and sequence diversity (Table 6). It achieved a sequence recovery of **78.03%**, representing an 11–22% improvement over PepFlow variants, and produced sequences with substantially lower Likeness scores (− 14.31), indicating distributions that are closer to natural peptides. At the same time, BC-Design maintained the highest diversity (**26.36%**), suggesting that it proposes a broad range of viable peptide solutions rather than collapsing onto a small set of sequences. These results show that biochemistry-aware inverse design can generate peptide variants that are accurate, natural, and diverse, which is critical for exploring functional sequence space around a given receptor complex.

**Table 6:** Evaluation of inverse design methods on the fixed-backbone sequence design task from the PepMerge dataset.

| Model | Sequence Recovery (%) ↑ | Worst % ↑ | Likeness ↓ | Diversity ↑ |
| --- | --- | --- | --- | --- |
| ProteinMPNN | 53.28 | 45.99 | -8.42 | 15.33 |
| ESM-IF | 43.51 | 36.18 | -8.39 | 13.76 |
| PepFlow w/Bb+Seq | 56.40 | 46.59 | -8.26 | 23.38 |
| PepFlow w/Bb+Seq+Ang | 54.32 | 44.48 | -8.58 | 20.65 |
| BC-DESIGN | 78.03 | 57.85 | -14.31 | 26.36 |

#### Antibody design

Finally, we evaluated BC-Design on antibody redesign using the AbMPNN benchmark [56] with structures drawn from SAbDab [57] and processed following AntiFold [58]. We focused on CDRH3, the most variable and functionally critical complementarity-determining region. BC-Design achieved a CDRH3 recovery rate of **96%**, substantially higher than alternative inverse folding models, and produced sampled CDR loops with the lowest backbone RMSD relative to experimentally resolved structures (Table 7). This indicates that the model can propose sequence variants that preserve fine-grained structural features required for antigen binding. In contrast, the backbone-only variant achieved sequence recovery comparable to AbMPNN but exhibited much higher RMSD, underscoring that backbone coordinates alone are insufficient to recover the structural specificity of antibody loops. The improved structural fidelity of BC-Design thus directly reflects the contribution of biochemical properties to capturing subtle sequence–structure constraints in antibodies.

**Table 7:** Model performance on the SAbDab dataset. Sequence recovery is evaluated on the CDRH3 design task, and RMSD is computed on sampled CDR loops.

| Model | Sequence Recovery (%) | RMSD (Å) |
| --- | --- | --- |
| ProteinMPNN | 35 | 1.03 |
| ESM-IF1 | 43 | 1.01 |
| AbMPNN | 56 | 0.98 |
| AntiFold | 60 | 0.95 |
| BC-DESIGN | 96 | 0.94 |
| BC-DESIGN (Backbone Only) | 54 | 1.54 |

### 3.5 Stratified Validation

Because BC-Design’s performance can vary with protein size and structural context, we conducted a stratified evaluation on the CATH 4.2 test set to assess robustness across diverse protein characteristics. This analysis highlights the model’s strengths and limitations under different structural, biochemical, and sequence conditions.

#### Protein size

We first stratified proteins by length and measured sequence- and structure-level performance across size bins (Fig. 5(a–d)). BC-Design achieves consistently strong results across all groups, with median sequence recovery exceeding **80%**, nssr above **90%**, and structural metrics (Ref-TM and pLDDT) above **0.8**. Sequence-level accuracy decreases moderately for longer proteins, reflecting the increased complexity and redundancy present in larger folds. In contrast, structural metrics remain robust even in the [400,500) range, suggesting that the model prioritizes fold-level correctness over exact residue identities as protein size grows. This behavior is consistent with the presence of sequence variability in peripheral regions that exert limited influence on global fold.

**Fig. 5:**
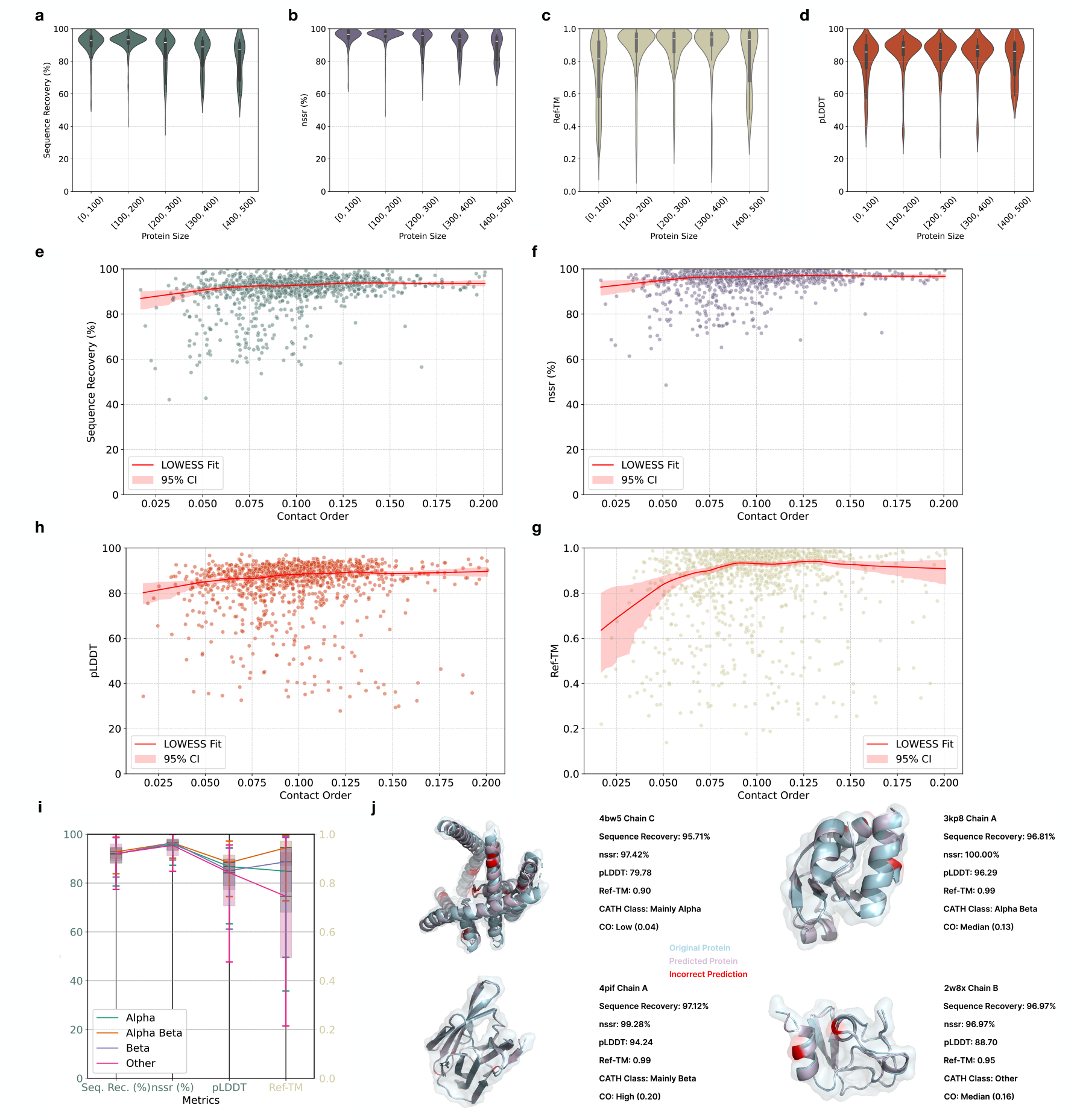
(**a–d**) Sequence- and structure-level performance of BC-Design across protein length bins in the CATH 4.2 test set. Median sequence recovery exceeds **80%**, nssr exceeds **85%**, and structural metrics (Ref-TM > 0.8, pLDDT*>* 0.75) remain high for all size ranges. Although recovery decreases slightly for larger proteins, structural fidelity is consistently maintained. **(e–h)** Trends with structural complexity, measured by contact order. All metrics show improving medians and reduced variance at higher contact orders, indicating that long-range interactions provide stronger geometric and biochemical constraints that facilitate accurate prediction. LOWESS curves highlight this positive dependence across metrics. **(i)** Performance across CATH classes. Sequence-level metrics remain stable across classes, whereas structural scores vary: Alpha–Beta proteins show the highest Ref-TM, while Beta and Other classes display greater variability, consistent with their more heterogeneous or beta-rich folds. **(j)** Representative case studies spanning four CATH classes and varying complexities. Across Mainly Alpha, Alpha–Beta, Mainly Beta, and Other folds, BC-Design achieves high recovery and structural accuracy, with highlighted regions illustrating both precise matches and occasional mismatches.

#### Structural complexity

To evaluate how folding topology affects model performance, we stratified proteins by contact order—a measure of the average sequence separation between contacting residues. As shown in Fig. 5(e–h), all four metrics exhibit higher medians and reduced spread as contact order increases. This indicates that proteins with more long-range interactions provide stronger geometrical and biochemical constraints that support accurate prediction. Despite these trends, performance differences across contact-order groups remain modest: case studies in Fig. 5(j) show that BC-Design maintains high sequence recovery and structural agreement across low-, medium-, and high-complexity proteins. Overall, the method is robust to variation in folding complexity.

#### CATH class

We next analyzed performance across CATH structural classes. Each protein was assigned the class of its largest domain, and the ‘Few Secondary Structures’ category was merged into an ‘Other’ group due to its small size. As shown in Fig. 5(i), sequence-level performance (sequence recovery and nssr) is uniformly high across all classes, demonstrating that BC-Design generalizes well across diverse secondary-structure compositions. Structural metrics, however, exhibit clear differences across classes. Kruskal–Wallis and Dunn’s post hoc tests show significant differences in pLDDT between the ‘Mainly Beta’ class and the ‘Mainly Alpha’ and ‘Alpha Beta’ classes, as well as differences in Ref-TM, with ‘Alpha Beta’ achieving the highest median. These patterns suggest that mixed *α*/*β* architectures—which combine stabilizing long-range *β*-sheet interactions with supportive *α*-helical packing—provide particularly well-constrained design contexts. Conversely, the ‘Other’ class, which includes irregular or weakly structured proteins, shows lower structural accuracy and greater variability, reflecting intrinsic modeling challenges. Representative examples for each class are shown in Fig. 5(j).

#### Protein region

To examine regional effects within proteins, we compared performance on core versus surface residues. As shown in Fig. 4(a), both sequence recovery and nssr are substantially higher in the core, consistent with the greater evolutionary conservation and structural determinism of buried positions. Surface residues—which often lie in flexible loops or solvent-exposed regions—show lower accuracy, mirroring prior observations in structure-only inverse folding. These results confirm that BC-Design more effectively captures stabilizing features essential for maintaining fold integrity.

#### Amino acid type

Finally, we assessed prediction quality across amino acid types using accuracy, precision, recall, and F1-score (Fig. 4(b)). Accuracy remains high across all residues, but several amino acids, including D, E, M, N, and Q, exhibit lower precision and recall. This reduction can be attributed to their biochemical and structural contexts: (i) D and E share similar negative charges and hydrophilicity; N and Q are polar amides; and M is uncharged but surface-favoring, making these residues difficult to distinguish when biochemical properties are locally similar. (ii) These residues frequently occur in dynamic surface regions or loops, where structural cues are weaker and conformational heterogeneity is higher. The correspondence between residue-level and region-level results reinforces the interpretation that biochemical ambiguity and structural flexibility, rather than model limitations, underlie these reduced scores.

### 3.6 BC-Design for functionally improved sequence design

To illustrate the practical applicability of BC-Design, we embedded it within a complete inverse-design workflow that integrates ThermoMPNN [59], RFDiffusion [60], and Boltz2 [61] (Fig. 6(a)). As a case study, we redesigned the phosphoglycerate kinase (PGK) enzyme (PDB ID: 16PK) to increase its affinity for the bisubstrate analog adenylyl 1,1,5,5-tetrafluoropentane-1,5-bisphosphonate (BIS) relative to the wild type (WT) [62]. PGK catalyzes a key step in glycolysis, the sole energy source for *Trypanosoma brucei* [63], the causative agent of African sleeping sickness. Designing PGK variants with enhanced BIS binding therefore has direct translational relevance, and PGK’s length (over 400 residues) makes it a stringent test for our method.

**Fig. 6:**
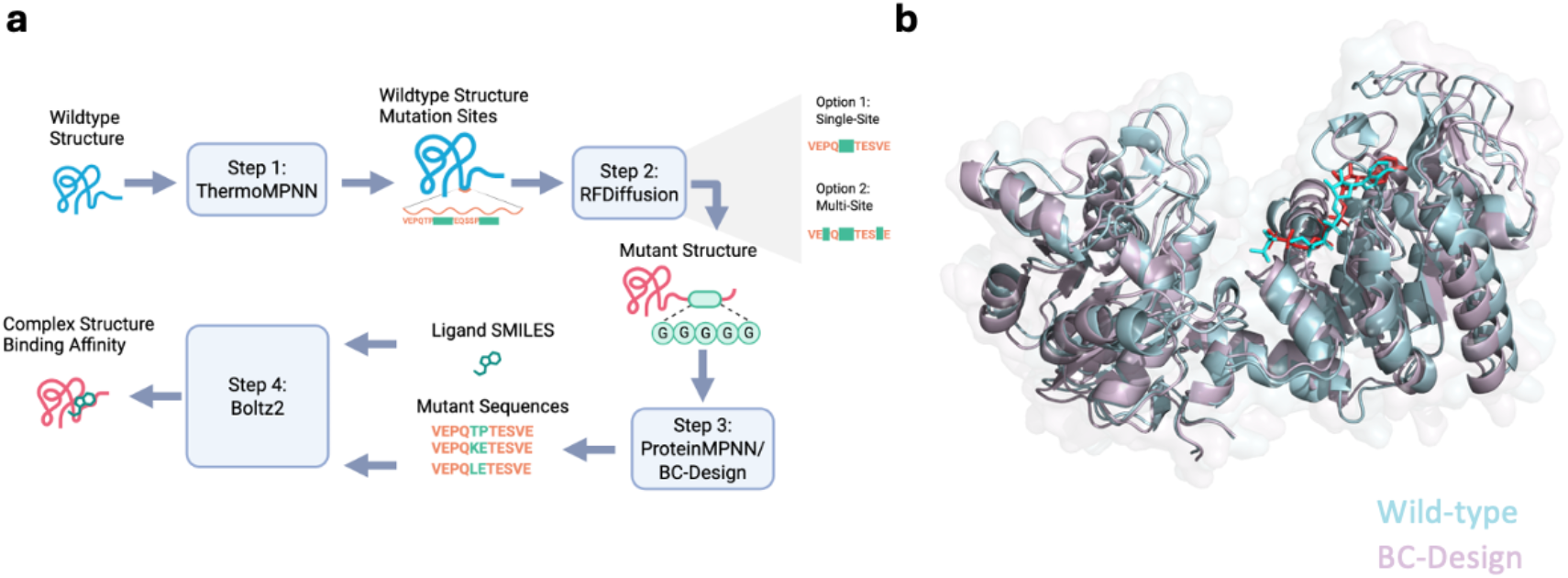
**(a)** Overview of the partial redesign workflow for PGK. Step 1: Wild-type PDB structures are input to ThermoMPNN to obtain residue-wise thermodynamic perturbation scores and identify candidate mutation regions. Step 2: The WT structure and selected regions are provided to RFDiffusion, which generates mutant backbones in which target residues are represented as glycine, corresponding to backbone-only geometry. We consider two prompting modes: a single-site prompt, where only one region is redesigned, and a multi-site prompt, where multiple regions are redesigned simultaneously. Step 3: The resulting mutant backbones are input to ProteinMPNN or BC-Design to recover full sequences. For BC-Design, biochemical features for non-mutated residues are provided to supply local physicochemical context. Step 4: Mutant sequences and ligand SMILES are input to Boltz2 to predict protein–ligand complex structures and estimate binding affinities. **(b)** PyMOL visualization of WT PGK (PDB ID: 16PK) and a BC-Design multi-site mutant in complex with BIS. Pale blue, WT PGK; cyan, ligand in the WT complex; pale purple, mutant PGK; red, ligand in the mutant complex.

We first passed the WT structure through ThermoMPNN to obtain residue-wise thermodynamic perturbation scores and identify regions where mutations are thermodynamically tolerated and potentially beneficial. RFDiffusion was then used to generate mutant backbones in these regions: in the single-site setting, only one region was specified, whereas in the multi-site setting, multiple regions were simultaneously targeted. In both cases, mutated positions were represented as glycine to reflect backbone-only information. The resulting partially redesigned backbones were then used for sequence recovery with either ProteinMPNN or BC-Design. When using BC-Design, biochemical features of the unmutated residues were supplied to provide a realistic physicochemical context for predicting mutated identities. Finally, we co-folded each mutant sequence together with the ligand SMILES using Boltz2 (Fig. 6(b)) and compared the predicted binding affinity to that of the WT. We use Boltz2 rather than AlphaFold2 because Boltz2 supports joint co-folding of protein–ligand complexes and provides direct estimates of ligand-binding energetics, whereas AF2 lacks native support for small-molecule cofactors and does not model protein–ligand interactions reliably.

Despite the considerable length of PGK, BC-Design produced mutants with substantially improved BIS affinity compared to both WT and a ProteinMPNN-designed variant. In the single-site setting, BC-Design yielded a 26.72% improvement in binding affinity, compared with 16.63% for ProteinMPNN. In the multi-site setting, BC-Design achieved a 51.02% improvement, versus 40.71% for ProteinMPNN. We also evaluated a biochemical-feature-unaware variant that uses only backbone coordinates as input. This backbone-only model produced smaller gains (24.14% and 31.45% in the single-site and multi-site settings, respectively), underscoring the added value of incorporating residue-level biochemical properties in guiding functional sequence optimization (Table 8).

**Table 8:** Binding-affinity improvements for PGK mutants designed by ProteinMPNN, a backbone-only variant of BC-Design, and the full biochemistry-aware BC-Design. Affinities are reported as percentage improvement in IC_50_ relative to the wild type. Single-site and Multi-sites correspond to the two prompting modes in Fig. 6.

| Model | ΔBinding Affinity (%) | |
| --- | --- | --- |
|  | Single-site | Multi-sites |
| ProteinMPNN | 16.63 | 40.71 |
| BC-DESIGN (Backbone Only) | 24.14 | 31.45 |
| BC-DESIGN | **26.72** | **51.02** |

### 3.7 Ablation Study on Staged Training and Contrastive Learning

To assess the contribution of the four-stage training curriculum, we compared BC-Design against an end-to-end variant in which all parameters were optimized jointly from scratch (Table 9).

**Table 9:** Comparison of BC-Design trained using the proposed four-stage curriculum versus an end-to-end from-scratch approach on the CATH 4.2 test set. Staged training substantially improves the backbone-only (BS) model, indicating a more robust structural encoder.

| Training Strategy | Sequence Recovery (%) | Perplexity | nssr (%) | Ref-TM | pLDDT |
| --- | --- | --- | --- | --- | --- |
| BC-DESIGN (Staged Training) | 90.25 | 1.38 | 94.71 | 0.85 | 83.79 |
| BC-DESIGN (From Scratch) | 91.30 | 1.31 | 95.62 | 0.85 | 84.04 |
| BC-DESIGN-BS (Staged Training) | 49.23 | 4.79 | 66.78 | 0.81 | 80.22 |
| BC-DESIGN-BS (From Scratch) | 45.87 | 5.08 | 64.39 | 0.80 | 78.83 |

**Table 10:**
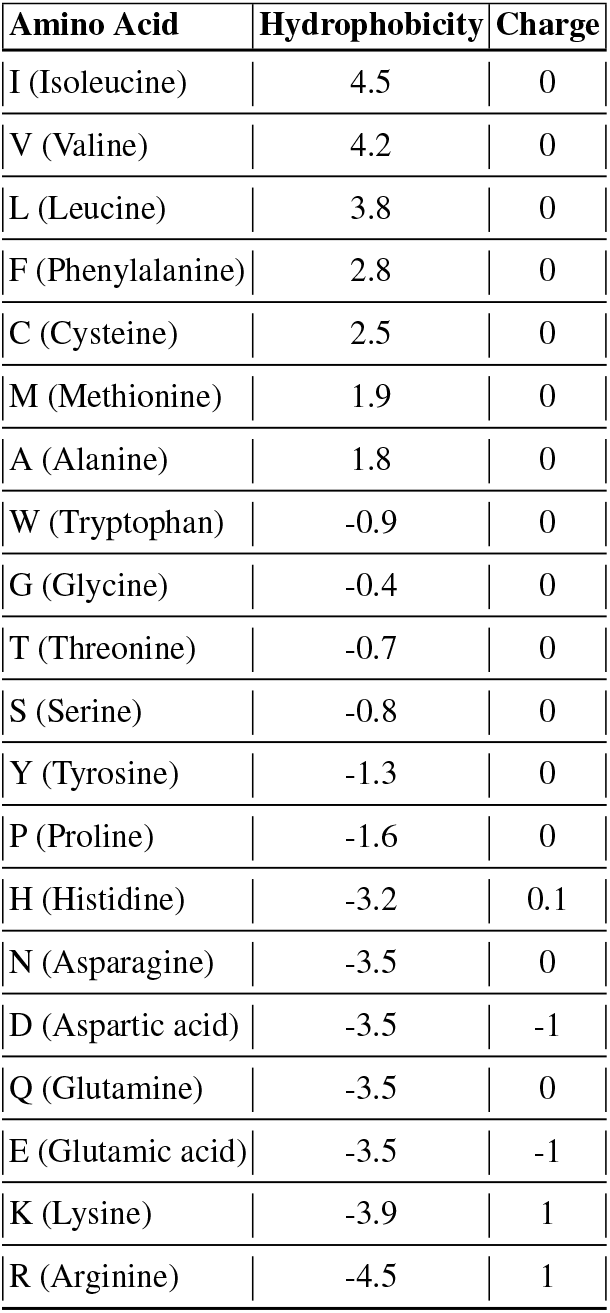
Hydrophobicity and charge values of amino acids.

| Amino Acid | Hydrophobicity | Charge |
| --- | --- | --- |
| I (Isoleucine) | 4.5 | 0 |
| V (Valine) | 4.2 | 0 |
| L (Leucine) | 3.8 | 0 |
| F (Phenylalanine) | 2.8 | 0 |
| C (Cysteine) | 2.5 | 0 |
| M (Methionine) | 1.9 | 0 |
| A (Alanine) | 1.8 | 0 |
| W (Tryptophan) | -0.9 | 0 |
| G (Glycine) | -0.4 | 0 |
| T (Threonine) | -0.7 | 0 |
| S (Serine) | -0.8 | 0 |
| Y (Tyrosine) | -1.3 | 0 |
| P (Proline) | -1.6 | 0 |
| H (Histidine) | -3.2 | 0.1 |
| N (Asparagine) | -3.5 | 0 |
| D (Aspartic acid) | -3.5 | -1 |
| Q (Glutamine) | -3.5 | 0 |
| E (Glutamic acid) | -3.5 | -1 |
| K (Lysine) | -3.9 | 1 |
| R (Arginine) | -4.5 | 1 |

When full biochemical and structural information is provided, both models achieve similar performance, with the end-to-end variant showing a marginally higher sequence recovery (91.30% vs. 90.25%). However, this advantage disappears in reduced-information settings. Under backbone-only inference, the staged-training model performs substantially better (49.23% vs. 45.87%), indicating that the structural encoder learned through the staged curriculum is more robust and self-sufficient.

These results highlight the primary benefit of the staged approach. By pretraining the structural encoder separately and aligning it with biochemical representations through global and local contrastive objectives, the model learns separated yet complementary modality-specific features. In contrast, end-to-end training encourages co-adaptation, yielding representations that rely more heavily on explicit biochemical cues and degrade more sharply when those cues are removed.

## 4 Discussion

Inverse protein folding is commonly evaluated using sequence recovery, yet the relationship between recovery, fold specification, and design utility is more nuanced than any single metric can convey. Our findings show that continuous biochemical properties offer a principled way to couple geometric structure with physicochemical environment, enabling the model to operate across a range of design regimes rather than committing to a single interpretation of the inverse folding objective. Below, we synthesize the conceptual implications of this biochemistry-aware formulation and outline its relevance for functional protein design.

### 4.1 Rethinking Sequence Recovery in the Context of Fold Identity and Design Objectives

Classical studies in structural biology have long established that moderate sequence identity—often as low as 30– 40%—is sufficient to determine a protein’s fold. This places natural bounds on the interpretability of high recovery values and highlights that replicating the wild-type sequence is not synonymous with recovering the biophysical constraints that underlie foldability. In this light, BC-Design should be understood not as a mechanism for reproducing native sequences, but as a model that learns how geometry and biochemical environment jointly constrain viable sequences.

The ability to modulate recovery through controlled masking of biochemical features further illustrates this distinction. High recovery represents a conservative operating point appropriate for stability-preserving redesign, while reduced recovery corresponds to more exploratory mutational behavior. Sequence recovery is therefore not the primary target of the model, but one coordinate in a tunable design landscape that can be adapted to the goals of a specific engineering task.

### 4.2 Biochemistry-aware Modeling Without Sequence Leakage

Incorporating biochemical information into inverse folding raises the question of whether such signals might inadvertently encode residue identity. Hydrophobicity and charge, however, are inherently degenerate properties shared across amino acid classes. When represented as spatially distributed properties on point-cloud samples, these features become continuous environmental descriptors rather than residue-specific labels.

The strong dependence of accuracy on the presence of these properties—manifested by a substantial drop under full masking—demonstrates that the model treats biochemical cues as contextual signals rather than shortcuts. Generalization analyses offer further evidence: performance on structurally large proteins declines when such proteins are underrepresented in the training set, and improves immediately when additional long-chain examples are introduced. This behavior aligns with genuine structure–biochemistry learning rather than implicit sequence encoding or memorization.

### 4.3 Controllable Fidelity–diversity Trade-offs for Protein Engineering

Systematic masking of biochemical features reveals a smooth, continuous frontier linking sequence fidelity and diversity. Full biochemical context yields high-fidelity designs, whereas full masking produces broad sequence exploration. This tunable range enables explicit and intuitive control over design behavior: users may adopt high-fidelity settings for conservative redesign, or high-diversity settings for tasks such as interface rewiring, specificity shifts, or binder evolution.

Notably, structural metrics remain stable across this spectrum, indicating that geometric fidelity can be preserved even as sequence diversity increases. This property allows the model to explore alternative solutions while maintaining foldability—an essential capability for functional protein engineering.

### 4.4 Implications for Function-oriented and Generative Protein Design

Across enzyme redesign, peptide–receptor engineering, and antibody CDR modeling, biochemistry-aware inverse folding improves functional metrics beyond what purely structure-based methods can achieve. This demonstrates that continuous physicochemical properties capture aspects of the energetic landscape that are directly relevant to functional behavior yet absent from backbone-only representations.

Beyond these task-specific gains, BC-Design provides a flexible foundation for generative protein design. Its ability to interpolate between high-fidelity and exploratory regimes, together with its robustness to structural noise and size variation, supports a wide range of applications—from conservative scaffold optimization to de novo functional exploration. Integrating geometric and biochemical information in a unified representation therefore offers a scalable strategy for modeling the chemical determinants of protein behavior, and points toward future protein design frameworks in which function, rather than native-sequence recovery, defines the primary objective.

Finally, the modular treatment of biochemical features affords practical flexibility during inference. Because physicochemical properties are supplied as an optional, spatially localized input channel, the model naturally supports region-specific masking: users can retain biochemical context in conserved scaffold regions while withholding it in targeted functional sites where sequence exploration is desired. This inference-time control reflects how protein designers typically operate and is independent of the contrastive or staged training strategy itself.

## 5 Conclusion

Our biochemistry-aware framework enables the model to operate across a range of design regimes, from high-fidelity sequence reconstruction to broad exploratory diversification, controlled through a simple masking mechanism that modulates the contribution of biochemical context. Across enzyme engineering, peptide–receptor design, and antibody CDR modeling, integrating structural geometry with continuous physicochemical properties consistently improves functional performance beyond what structure-only models can achieve. The model’s behavior under biochemical masking, out-of-distribution conditions, and noisy backbone inputs further demonstrates that it learns genuine structure–biochemistry relationships rather than memorizing sequence patterns, providing explicit and interpretable control over the fidelity–diversity trade-off central to practical protein design. Looking forward, the framework naturally extends to settings with predicted or uncertain structures and to integration with de novo backbone generation. We anticipate that such models will form a core component of next-generation, function-oriented protein design pipelines.

## A Comparison of Biochemical Feature Representations

In this section, we compare our biochemical feature representation with existing approaches which largely fall into two categories.

### Per-residue biochemical annotation

A common strategy is to treat biochemical properties as discrete attributes attached to each residue node in a residue-level protein graph. In this formulation, the protein is represented as a graph whose nodes correspond to amino acids and whose edges capture geometric relationships. Biochemical features are introduced as additional node features. While straightforward and computationally efficient, this representation is inherently localized: the biochemical properties of a residue are only defined at the residue center and do not reflect the continuous spatial variation of physicochemical fields across the protein surface or interior. As a result, this approach cannot model how biochemical environments extend beyond residue coordinates.

### Surface-based biochemical annotation

A second line of work constructs an explicit molecular surface, typically via solvent-accessible or solvent-excluded surface algorithms, and assigns biochemical properties to points on this surface. This representation captures the chemical geometry experienced by interacting partners, making it well suited for binder, peptide, and interface design. However, because biochemical features are restricted to surface points, they fail to encode the physicochemical context of internal cavities, buried residues, or broader volumetric environments that strongly influence folding stability and global energetics. Consequently, surface-based approaches represent only the exterior chemical landscape and cannot model the full 3D distribution of biochemical constraints.

### Our volumetric, continuous biochemistry representation

BC-Design introduces a distinct representation that departs from both residue-centric and surface-centric formulations. We model biochemical properties as continuous spatial distributions defined throughout the entire 3D volume occupied by the protein. By sampling a unified point cloud spanning both the molecular surface and interior regions, and assigning hydrophobicity and charge as biochemical signals, this representation captures the global physicochemical landscape that governs folding. This volumetric perspective allows the model to reason about biochemical gradients and long-range physicochemical context, enabling a principled and flexible integration of structure and biochemistry.

A visual summary of these three representations is provided in Fig. 7, illustrating the conceptual differences and highlighting the unique expressiveness of our approach.

**Fig. 7:**
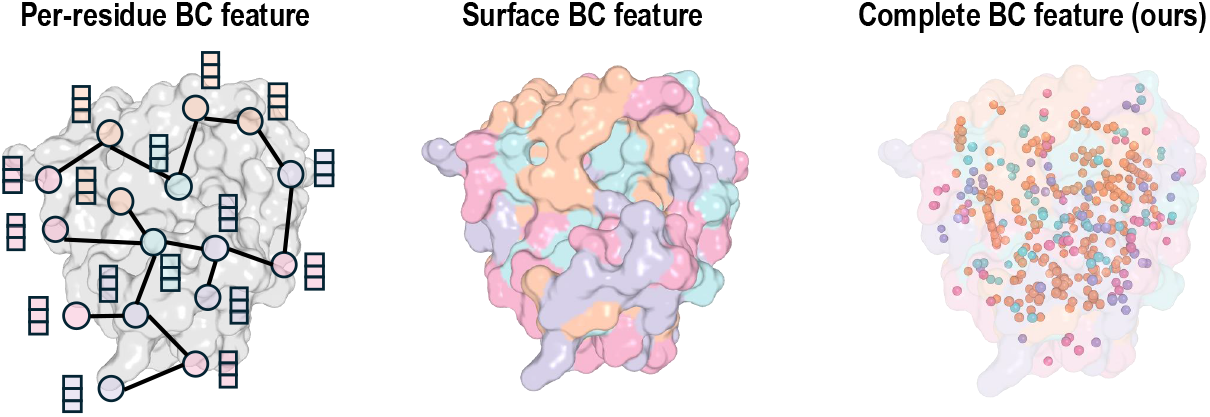
Illustration of three biochemical (BC) feature representations. Per-residue approaches encode biochemical properties only at residue nodes in a protein graph. Surface-based methods assign properties to points on the molecular surface. BC-Design introduces a volumetric, continuous representation that distributes biochemical signals throughout the entire 3D protein volume, including both surface and interior regions.

## B Biochemical Features

Here, we present the biochemical properties of all 20 standard amino acids, specifically their hydrophobicity and charge values. These values form the foundation for generating biochemical feature distributions in our work. The hydrophobicity values are derived from the Kyte-Doolittle Hydrophobicity Scale obtained from the ImMunoGeneTics information system.

In this table, hydrophobicity values range from −4.5 to 4.5, where: Positive values indicate hydrophobic amino acids (e.g., Isoleucine: 4.5, Valine: 4.2, Leucine: 3.8). Negative values indicate hydrophilic amino acids (e.g., Arginine: −4.5, Lysine: −3.9).

The charge values are simplified to represent the ionic state at physiological pH: +1 for positively charged amino acids (Lysine, Arginine). −1 for negatively charged amino acids (Aspartic acid, Glutamic acid) 0 for neutral amino acids. 0.1 for Histidine due to its special properties at physiological pH.

These biochemical features play a crucial role in our BC-Design framework, as they help capture the physical and chemical properties that determine protein folding and stability. By incorporating both hydrophobicity and charge distributions, our model can better understand and predict the amino acid sequences that would adopt a desired protein structure.

## C Fairness of Biochemical Feature Introduction

The introduction of biochemical features in our model raises potential concerns about information leakage, particularly whether these features could unfairly encode sequence information. Here we demonstrate the fairness of our approach from two key aspects:

### C.1 Universality of Biochemical Properties

The biochemical features (hydrophobicity and charge) assigned to point clouds represent universal physicochemical properties of proteins rather than sequence-specific information. Multiple amino acids share similar biochemical characteristics, making it impossible to uniquely determine residue types based on these features alone:

1. Hydrophobicity groups:
  - Highly hydrophobic: Isoleucine (4.5), Valine (4.2), Leucine (3.8)
  - Moderately hydrophobic: Phenylalanine (2.8), Cysteine (2.5)
  - Highly hydrophilic: Lysine (− 3.9), Arginine (− 4.5)
  - Moderately hydrophilic: Asparagine (− 3.5), Glutamine (− 3.5), Aspartic acid (− 3.5), Glutamic acid (− 3.5)
2. Charge groups:
  - Positively charged (+1): Lysine, Arginine
  - Negatively charged (− 1): Aspartic acid, Glutamic acid
  - Neutral (0): Majority of amino acids

This degeneracy in biochemical properties means that multiple amino acid sequences could potentially satisfy the same biochemical feature distribution, making it impossible to reconstruct the original sequence solely from these features.

### C.2 Randomness in Point Cloud Generation

The point cloud generation process incorporates several layers of randomness that further prevent sequence information leakage:

1. Spatial sampling:
  - Points are uniformly sampled within the bounding box
  - The final point cloud is randomly subsampled to 5000 points
2. Feature assignment: The continuous and random nature of the point cloud representation blurs discrete residue positions

These characteristics ensure that while the biochemical features provide valuable information about the chemical environment necessary for protein folding, they do not encode or leak sequence information that would make the inverse folding task trivial. This maintains the fundamental challenge of the task while enriching the model’s understanding of the physicochemical constraints that guide protein folding.

## D Implementation Details

Our BC-Design architecture employs a hidden size of 256, featuring a Struct-Encoder with 3 Structure Graph Transformer Layers and 8 attention heads, complemented by a BC-Encoder with 4 hierarchical levels and a 4-head MHA block. The fusion component incorporates 3 BC-Fusion blocks, each equipped with 8 attention heads. We train the model for 20 epochs on NVIDIA A100 GPUs using the AdamW optimizer and OneCycle learning rate scheduler, with a batch size of 4 and a learning rate of 0.0002. The model training requires a total of 6h 19m for 20 epochs on 4 NVIDIA A100 GPUs. During inference, evaluated on a single A100 GPU with a batch size of 1, the model achieves an average inference time of 0.0651 seconds per batch, with a standard deviation of 0.1450 seconds.

## E Future Work

Building on our success, several promising research directions could further enhance the approach:

### E.1 Experimental Validation through Iterative Design Cycles

Unlike purely language model-based approaches, implementing a Trust Region-based optimization approach with experimental feedback loops could significantly improve real-world applicability. This would involve designing initial peptides, obtaining experimental effectiveness data, and refining designs through multiple iterations to achieve optimal performance against specific targets like fungi. The iterative nature of this approach allows for continuous improvement based on real experimental outcomes, potentially leading to more biologically relevant designs than those created through computational methods alone.

### E.2 User-specified Biochemical Property Distributions

Currently, BC-Design derives biochemical features from ground-truth sequences during evaluation, which may not fully reflect real-world design scenarios. Future work could explore methods for protein design with user-specified biochemical property distributions, effectively decoupling feature generation from sequence recovery. This advancement would enable designers to directly control the physicochemical environment of the protein, specifying desired hydrophobicity and charge distributions without relying on existing sequences. Such capability would significantly expand the creative possibilities for protein engineers, allowing them to explore novel sequence spaces that satisfy specific biochemical criteria while maintaining structural integrity.

By addressing these areas, BC-Design could evolve into an even more powerful tool for computational protein engineering and therapeutic development, bridging the gap between computational design and experimental validation through a more iterative and constraint-aware approach.

## F Comparison Between GVP-GNN and BC-Design Approaches

GVP-GNN [30] and BC-Design represent two significant advancements in learning from protein structure, yet they differ in several fundamental aspects. GVP-GNN primarily focuses on augmenting graph neural networks with geometric vector perceptrons to perform geometric reasoning while maintaining equivariance. It operates on both scalar and vector features, enabling direct representation of 3D information throughout graph propagation without reducing such information to scalars that may not fully capture complex geometry. The model learns to encode the 3D geometry through vector representations that transform appropriately under spatial rotations, creating a global coordinate system across the structure.

BC-Design, in contrast, introduces an approach that explicitly represents biochemical properties as continuous distributions throughout the protein structure. Rather than encoding biochemical features as discrete properties of individual residues, it models hydrophobicity and charge as decorations on randomly sampled points across both exterior surfaces and internally bound regions. This provides a more natural way to capture the spatial distribution of biochemical properties as they exist in folded protein states, moving beyond the discrete residue-level representation prevalent in traditional models.

The architectural design of these two approaches also differs significantly. GVP-GNN employs relatively simple geometric vector perceptrons as its core computational unit, where vector channels directly encode geometric features. BC-Design implements a more complex architecture comprising a STRUCT-ENCODER that processes residue-level structural information through a hierarchical graph transformer, a BC-ENCODER that handles biochemical features, and a BC-FUSION module that integrates structure and biochemistry through a bipartite graph structure. This multi-component design allows BC-Design to process and fuse multiple information modalities.

A key innovation in BC-Design not present in GVP-GNN is its use of contrastive learning to bridge training and inference phases. During training, BC-Design leverages both structural information and biochemical feature distributions to learn rich, biochemistry-aware embeddings. However, at inference time, it requires only the backbone structure as input, with the structural embeddings having implicitly captured biochemical information. This creates a model that harnesses biochemical awareness during training while maintaining practical simplicity during application.

Performance-wise, BC-Design demonstrates significant improvements over existing methods, including GVP-GNN, particularly in inverse protein folding tasks. BC-Design achieves 90% sequence recovery compared to state-of-the-art methods’ 67%, representing a 21% absolute improvement, and reduces perplexity from 2.4 to 1.38. These improvements stem from the model’s ability to capture and leverage biochemical context in protein sequence design, highlighting the importance of representing biochemical properties as continuous distributions rather than discrete features.

Both approaches maintain rotation invariance in their scalar outputs and equivariance in their vector outputs with respect to 3D transformations, an essential property for learning from protein structure. However, BC-Design’s biochemistry-aware approach appears to provide a more comprehensive framework for capturing the physical and chemical principles that govern protein folding and stability, leading to its superior performance on inverse protein folding tasks.

## G Discussion on Atom-Based Sampling for Biochemical Feature Construction

The current BC-Design approach represents biochemical features through point clouds that sample both protein surfaces and internal spaces, demonstrating significant improvements in inverse protein folding. However, the uniform sampling strategy within a bounding box may not optimally capture the complex biochemical environment within proteins. Here, we explore the potential benefits and challenges of implementing an atom-based sampling approach as an alternative to the current methodology.

Atom-based sampling would leverage the precise atomic coordinates from the protein structure itself, rather than relying on geometric approximations. This approach could provide a more biologically accurate representation of the biochemical environment by directly sampling points based on actual atomic positions. Such sampling would inherently capture the non-uniform distribution of biochemical properties throughout the protein volume, potentially improving the model’s ability to learn subtle patterns that influence amino acid preferences at specific positions. Furthermore, by correlating sampling density with functionally important regions or atoms, the model could develop enhanced sensitivity to critical areas that determine protein function.

A key advantage of atom-based sampling would be its ability to better represent the discrete nature of protein biochemistry while maintaining our continuous distribution paradigm. Different atom types contribute distinctly to the overall biochemical environment—polar atoms create hydrophilic regions, while carbon-rich areas form hydrophobic clusters. By basing our sampling on these atomic coordinates and types, we could more accurately represent the gradients and boundaries between different biochemical zones within the protein structure. This could be particularly beneficial for capturing features like binding pockets, catalytic sites, or stabilizing hydrophobic cores.

Implementation of atom-based sampling would require careful consideration of several factors. First, proteins contain varying numbers of atoms, necessitating adaptive sampling techniques to maintain consistent representation sizes across different structures. Second, a weighting scheme would need to be developed to determine sampling density around different atom types, potentially giving preference to side chain atoms that more strongly influence biochemical properties. Finally, computational efficiency must be considered, as the increased complexity of atom-based sampling could impact training and inference times.

Despite these challenges, atom-based sampling presents a promising direction for enhancing the biochemical awareness of our model. The potential improvements in capturing atomic-level biochemical environments could further refine sequence predictions, particularly for functionally specialized regions where precise biochemical conditions are essential for protein activity. Future work will explore hybrid approaches that combine the computational efficiency of uniform sampling with the biochemical accuracy of atom-based methods, potentially leading to even more accurate and biologically relevant protein designs.

## Bibliography

[1] Zhangyang Gao, Cheng Tan, Yijie Zhang, Xingran Chen, Lirong Wu, and Stan Z Li. Proteininvbench: Benchmarking protein inverse folding on diverse tasks, models, and metrics. Advances in Neural Information Processing Systems, 36, 2024.

[2] Zhangyang Gao, Cheng Tan, Pablo Chacón, and Stan Z Li. Pifold: Toward effective and efficient protein inverse folding. arXiv preprint arXiv:2209.12643, 2022.

[3] Matthew McPartlon and Jinbo Xu. An end-to-end deep learning method for protein side-chain packing and inverse folding. Proceedings of the National Academy of Sciences, 120(23):e2216438120, 2023.

[4] Chloe Hsu, Robert Verkuil, Jason Liu, Zeming Lin, Brian Hie, Tom Sercu, Adam Lerer, and Alexander Rives. Learning inverse folding from millions of predicted structures. In International conference on machine learning, pages 8946–8970. PMLR, 2022.

[5] Natalie Maus, Yimeng Zeng, Daniel Allen Anderson, Phillip Maffettone, Aaron Solomon, Peyton Greenside, Osbert Bastani, and Jacob R Gardner. Inverse protein folding using deep bayesian optimization. arXiv preprint arXiv:2305.18089, 2023.

[6] Rohith Krishna, Jue Wang, Woody Ahern, Pascal Sturmfels, Preetham Venkatesh, Indrek Kalvet, Gyu Rie Lee, Felix S Morey-Burrows, Ivan Anishchenko, Ian R Humphreys, et al. Generalized biomolecular modeling and design with rosettafold all-atom. Science, 384(6693):eadl2528, 2024.

[7] Bowen Jing, Stephan Eismann, Pratham N Soni, and Ron O Dror. Equivariant graph neural networks for 3d macromolecular structure. arXiv preprint arXiv:2106.03843, 2021.

[8] John Jumper, Richard Evans, Alexander Pritzel, Tim Green, Michael Figurnov, Olaf Ronneberger, Kathryn Tunyasuvunakool, Russ Bates, Augustin Žídek, Anna Potapenko, et al. Highly accurate protein structure prediction with alphafold. nature, 596(7873): 583–589, 2021.

[9] Julia Koehler Leman, Pawel Szczerbiak, P Douglas Renfrew, Vladimir Gligorijevic, Daniel Berenberg, Tommi Vatanen, Bryn C Taylor, Chris Chandler, Stefan Janssen, Andras Pataki, et al. Sequence-structure-function relationships in the microbial protein universe. Nature communications, 14(1): 2351, 2023.

[10] Varun R Shanker, Theodora UJ Bruun, Brian L Hie, and Peter S Kim. Inverse folding of protein complexes with a structure-informed language model enables unsupervised antibody evolution. bioRxiv, 2023.

[11] Daniel Cutting, Frédéric A Dreyer, David Errington, Constantin Schneider, and Charlotte M Deane. De novo antibody design with se (3) diffusion. arXiv preprint arXiv:2405.07622, 2024.

[12] Pascal Notin, Nathan Rollins, Yarin Gal, Chris Sander, and Debora Marks. Machine learning for functional protein design. Nature biotechnology, 42(2): 216–228, 2024.

[13] Zhixiu Li, Yuedong Yang, Eshel Faraggi, Jian Zhan, and Yaoqi Zhou. Direct prediction of profiles of sequences compatible with a protein structure by neural networks with fragment-based local and energy-based nonlocal profiles. Proteins: Structure, Function, and Bioinformatics, 82(10): 2565–2573, 2014.

[14] James O’Connell, Zhixiu Li, Jack Hanson, Rhys Heffernan, James Lyons, Kuldip Paliwal, Abdollah Dehzangi, Yuedong Yang, and Yaoqi Zhou. Spin2: Predicting sequence profiles from protein structures using deep neural networks. Proteins: Structure, Function, and Bioinformatics, 86(6): 629–633, 2018.

[15] Yuan Zhang, Yang Chen, Chenran Wang, Chun-Chao Lo, Xiuwen Liu, Wu Wei, and Jinfeng Zhang. Prodconnprotein design using a convolutional neural network. Biophysical Journal, 118(3):43a–44a, 2020.

[16] Yifei Qi and John ZH Zhang. Densecpd: improving the accuracy of neural-network-based computational protein sequence design with densenet. Journal of chemical information and modeling, 60(3): 1245–1252, 2020.

[17] Alexey Strokach, David Becerra, Carles Corbi-Verge, Albert Perez-Riba, and Philip M Kim. Fast and flexible protein design using deep graph neural networks. Cell systems, 11(4): 402–411, 2020.

[18] John Ingraham, Vikas Garg, Regina Barzilay, and Tommi Jaakkola. Generative models for graph-based protein design. Advances in neural information processing systems, 32, 2019.

[19] Justas Dauparas, Ivan Anishchenko, Nathaniel Bennett, Hua Bai, Robert J Ragotte, Lukas F Milles, Basile IM Wicky, Alexis Courbet, Rob J de Haas, Neville Bethel, et al. Robust deep learning–based protein sequence design using proteinmpnn. Science, 378(6615): 49–56, 2022.

[20] Nicolas Renaud, Cunliang Geng, Sonja Georgievska, Francesco Ambrosetti, Lars Ridder, Dario F Marzella, Manon F Réau, Alexandre MJJ Bonvin, and Li C Xue. Deeprank: a deep learning framework for data mining 3d protein-protein interfaces. Nature communications, 12(1): 7068, 2021.

[21] Yipin Lei, Shuya Li, Ziyi Liu, Fangping Wan, Tingzhong Tian, Shao Li, Dan Zhao, and Jianyang Zeng. A deeplearning framework for multi-level peptide–protein interaction prediction. Nature communications, 12(1): 5465, 2021.

[22] Raktim Mitra, Jinsen Li, Jared M Sagendorf, Yibei Jiang, Ari S Cohen, Tsu-Pei Chiu, Cameron J Glasscock, and Remo Rohs. Geometric deep learning of protein–dna binding specificity. Nature Methods, pages 1–10, 2024.

[23] Yanting Li, Jiyue Jiang, Zikang Wang, Ziqian Lin, Dongchen He, Yuheng Shan, Yanruisheng Shao, Jiayi Li, Xiangyu Shi, Jiuming Wang, et al. Ds-progen: A dual-structure deep language model for functional protein design. arXiv preprint arXiv:2505.12511, 2025.

[24] Fang Wu, Zhengyuan Zhou, Shuting Jin, Xiangxiang Zeng, Jure Leskovec, and Jinbo Xu. Surface-based molecular design with multi-modal flow matching. In Proceedings of the 31st ACM SIGKDD Conference on Knowledge Discovery and Data Mining V. 2, pages 3192–3203, 2025.

[25] Guanlue Li, Xufeng Zhao, Fang Wu, and Sören Laue. Joint design of protein surface and backbone using a diffusion bridge model. In The Thirty-ninth Annual Conference on Neural Information Processing Systems, 2025.

[26] Fang Wu, Shuting Jin, Jianmin Wang, Zerui Xu, Jinbo Xu, Brian Hie, et al. Surfdesign: Effective protein design on molecular surfaces.

[27] Zhangyang Gao, Jue Wang, Cheng Tan, Lirong Wu, Yufei Huang, Siyuan Li, Zhirui Ye, and Stan Z Li. Uniif: Unified molecule inverse folding. Advances in Neural Information Processing Systems, 37: 135843–135860, 2024.

[28] Zhenqiao Song, Tinglin Huang, Lei Li, and Wengong Jin. Surfpro: Functional protein design based on continuous surface. In Forty-first International Conference on Machine Learning.

[29] Christine A Orengo, Alex D Michie, Susan Jones, David T Jones, Mark B Swindells, and Janet M Thornton. Cath–a hierarchic classification of protein domain structures. Structure, 5(8): 1093–1109, 1997.

[30] Bowen Jing, Stephan Eismann, Patricia Suriana, Raphael JL Townshend, and Ron Dror. Learning from protein structure with geometric vector perceptrons. arXiv preprint arXiv:2009.01411, 2020.

[31] Cheng Tan, Zhangyang Gao, Jun Xia, Bozhen Hu, and Stan Z Li. Generative de novo protein design with global context. arXiv preprint arXiv:2204.10673, 2022.

[32] Zhangyang Gao, Cheng Tan, and Stan Z Li. Alphadesign: A graph protein design method and benchmark on alphafolddb. arXiv preprint arXiv:2202.01079, 2022.

[33] Xing Zhang, Hongmei Yin, Fei Ling, Jian Zhan, and Yaoqi Zhou. Spin-cgnn: Improved fixed backbone protein design with contact map-based graph construction and contact graph neural network. PLoS Computational Biology, 19(12):e1011330, 2023.

[34] Kai Yi, Bingxin Zhou, Yiqing Shen, Pietro Lio, and Yuguang Wang. Graph denoising diffusion for inverse protein folding. Advances in Neural Information Processing Systems, 36, 2024.

[35] Jieling He, Wenxu Wu, and Xiaowo Wang. Diprot: A deep learning based interactive toolkit for efficient and effective protein design. Synthetic and Systems Biotechnology, 9(2): 217–222, 2024.

[36] Hui Wang, Dong Liu, Kailong Zhao, Yajun Wang, and Guijun Zhang. Spdesign: protein sequence designer based on structural sequence profile using ultrafast shape recognition. Briefings in Bioinformatics, 25(3):bbae146, 2024.

[37] Xinyi Zhou, Guangyong Chen, Junjie Ye, Ercheng Wang, Jun Zhang, Cong Mao, Zhanwei Li, Jianye Hao, Xingxu Huang, Jin Tang, et al. Prorefiner: an entropy-based refining strategy for inverse protein folding with global graph attention. Nature Communications, 14(1): 7434, 2023.

[38] Zaixiang Zheng, Yifan Deng, Dongyu Xue, Yi Zhou, Fei Ye, and Quanquan Gu. Structure-informed language models are protein designers. In International conference on machine learning, pages 42317–42338. PMLR, 2023.

[39] Zhangyang Gao, Cheng Tan, and Stan Z Li. Knowledge-design: Pushing the limit of protein design via knowledge refinement. arXiv preprint arXiv:2305.15151, 2023.

[40] Jiangbin Zheng and Stan Z Li. Progressive multi-modality learning for inverse protein folding. In 2024 IEEE International Conference on Multimedia and Expo (ICME), pages 1–6. IEEE, 2024.

[41] Weian Mao, Muzhi Zhu, Zheng Sun, Shuaike Shen, Lin Yuanbo Wu, Hao Chen, and Chunhua Shen. De novo protein design using geometric vector field networks. arXiv preprint arXiv:2310.11802, 2023.

[42] Robert Verkuil, Ori Kabeli, Yilun Du, Basile IM Wicky, Lukas F Milles, Justas Dauparas, David Baker, Sergey Ovchinnikov, Tom Sercu, and Alexander Rives. Language models generalize beyond natural proteins. BioRxiv, pages 2022–12, 2022.

[43] Jue Wang, Sidney Lisanza, David Juergens, Doug Tischer, Joseph L Watson, Karla M Castro, Robert Ragotte, Amijai Saragovi, Lukas F Milles, Minkyung Baek, et al. Scaffolding protein functional sites using deep learning. Science, 377(6604): 387–394, 2022.

[44] Zeming Lin, Halil Akin, Roshan Rao, Brian Hie, Zhongkai Zhu, Wenting Lu, Allan dos Santos Costa, Maryam Fazel-Zarandi, Tom Sercu, Sal Candido, et al. Language models of protein sequences at the scale of evolution enable accurate structure prediction. BioRxiv, 2022:500902, 2022.

[45] Natalie L Dawson, Tony E Lewis, Sayoni Das, Jonathan G Lees, David Lee, Paul Ashford, Christine A Orengo, and Ian Sillitoe. Cath: an expanded resource to predict protein function through structure and sequence. Nucleic acids research, 45(D1):D289–D295, 2017.

[46] Patrick Löffler, Samuel Schmitz, Enrico Hupfeld, Reinhard Sterner, and Rainer Merkl. Rosetta: Msf: a modular framework for multi-state computational protein design. PLoS computational biology, 13(6):e1005600, 2017.

[47] Steven Henikoff and Jorja G Henikoff. Amino acid substitution matrices from protein blocks. Proceedings of the National Academy of Sciences, 89(22): 10915–10919, 1992.

[48] Zeming Lin, Halil Akin, Roshan Rao, Brian Hie, Zhongkai Zhu, Wenting Lu, Nikita Smetanin, Robert Verkuil, Ori Kabeli, Yaniv Shmueli, et al. Evolutionary-scale prediction of atomic-level protein structure with a language model. Science, 379(6637): 1123–1130, 2023.

[49] Chuanrui Wang, Bozitao Zhong, Zuobai Zhang, Narendra Chaudhary, Sanchit Misra, and Jian Tang. Pdb-struct: A comprehensive benchmark for structure-based protein design. arXiv preprint arXiv:2312.00080, 2023.

[50] Ruidong Wu, Fan Ding, Rui Wang, Rui Shen, Xiwen Zhang, Shitong Luo, Chenpeng Su, Zuofan Wu, Qi Xie, Bonnie Berger, et al. High-resolution de novo structure prediction from primary sequence. BioRxiv, pages 2022–07, 2022.

[51] Mukund Sundararajan, Ankur Taly, and Qiqi Yan. Axiomatic attribution for deep networks. In International conference on machine learning, pages 3319–3328. PMLR, 2017.

[52] Mihaly Varadi, Damian Bertoni, Paulyna Magana, Urmila Paramval, Ivanna Pidruchna, Malarvizhi Radhakrishnan, Maxim Tsenkov, Sreenath Nair, Milot Mirdita, Jingi Yeo, et al. Alphafold protein structure database in 2024: providing structure coverage for over 214 million protein sequences. Nucleic acids research, 52(D1):D368–D375, 2024.

[53] Alexander Kroll, Sahasra Ranjan, Martin KM Engqvist, and Martin J Lercher. A general model to predict small molecule substrates of enzymes based on machine and deep learning. Nature Communications, 14(1): 2787, 2023.

[54] Zhenqiao Song, Tinglin Huang, Lei Li, and Wengong Jin. Surfpro: Functional protein design based on continuous surface. arXiv preprint arXiv:2405.06693, 2024.

[55] Jiahan Li, Chaoran Cheng, Zuofan Wu, Ruihan Guo, Shitong Luo, Zhizhou Ren, Jian Peng, and Jianzhu Ma. Full-atom peptide design based on multi-modal flow matching. arXiv preprint arXiv:2406.00735, 2024.

[56] Frédéric A. Dreyer, Daniel Cutting, Constantin Schneider, Henry Kenlay, and Charlotte M. Deane. Inverse folding for antibody sequence design using deep learning, 2023.

[57] James Dunbar, Konrad Krawczyk, Jinwoo Leem, Terry Baker, Angelika Fuchs, Guy Georges, Jiye Shi, and Charlotte M. Deane. Sabdab: the structural antibody database. Nucleic Acids Research, 42(D1):D1140–D1146, 11 2013.

[58] Magnus Haraldson Høie, Alissa Hummer, Tobias H. Olsen, Broncio Aguilar-Sanjuan, Morten Nielsen, and Charlotte M. Deane. Antifold: Improved antibody structure-based design using inverse folding, 2024.

[59] Henry Dieckhaus, Michael Brocidiacono, Nicholas Z. Randolph, and Brian Kuhlman. Transfer learning to leverage larger datasets for improved prediction of protein stability changes. Proceedings of the National Academy of Sciences, 121(6), feb 2024.

[60] Joseph L. Watson, David Juergens, Nathaniel R. Bennett, Brian L. Trippe, Jason Yim, Helen E. Eisenach, Woody Ahern, Andrew J. Borst, Robert J. Ragotte, Lukas F. Milles, Basile I. M. Wicky, Nikita Hanikel, Samuel J. Pellock, Alexis Courbet, William Sheffler, Jue Wang, Preetham Venkatesh, Isaac Sappington, Susana Vázquez Torres, Anna Lauko, Valentin De Bortoli, Emile Mathieu, Sergey Ovchinnikov, Regina Barzilay, Tommi S. Jaakkola, Frank DiMaio, Minkyung Baek, and David Baker. De novo design of protein structure and function with RFdiffusion. Nature, 620(7976):1089–1100, aug 2023.

[61] Saro Passaro, Gabriele Corso, Jeremy Wohlwend, Mateo Reveiz, Stephan Thaler, Vignesh Ram Somnath, Noah Getz, Tally Portnoi, Julien Roy, Hannes Stark, David Kwabi-Addo, Dominique Beaini, Tommi Jaakkola, and Regina Barzilay. Boltz-2: Towards Accurate and Efficient Binding Affinity Prediction, jun 2025.

[62] Bradley E Bernstein, David M Williams, Jerome C Bressi, Peter Kuhn, Michael H Gelb, G.Michael Blackburn, and Wim G.J Hol. A bisubstrate analog induces unexpected conformational changes in phosphoglycerate kinase from trypanosoma brucei. Journal of Molecular Biology, 279:1137–1148, 6 1998.

[63] Onno Misset and Fred R. Opperdoes. The phosphoglycerate kinases from trypanosoma brucei. European Journal of Biochemistry, 162:493–500, 2 1987.

